# ReCom: A semi-supervised approach to ultra-tolerant database search for improved identification of modified peptides

**DOI:** 10.1101/2023.04.10.535358

**Authors:** Andrea Laguillo-Gómez, Enrique Calvo, Noa Martín-Cófreces, Marta Lozano-Prieto, Francisco Sánchez-Madrid, Jesús Vázquez

## Abstract

Open-search methods allow unbiased, high-throughput identification of post-translational modifications in proteins at an unprecedented scale. The performance of current open-search algorithms is diminished by experimental errors in the determination of the precursor peptide mass. In this work we propose a semi-supervised open search approach, called ReCom, that minimizes this effect by taking advantage of a priori known information from a reference database, such as Unimod or a database provided by the user. We present a proof-of-concept study using Comet-ReCom, an improved version of Comet-PTM. Comet-ReCom increased identification performance of Comet-PTM by 68%. This increased performance of Comet-ReCom to score the MS/MS spectrum comes in parallel with a significantly better assignation of the monoisotopic peak of the precursor peptide in the MS spectrum, even in cases of peptide coelution. Our data demonstrate that open searches using ultra-tolerant mass windows can benefit from using a semi-supervised approach that takes advantage from previous knowledge on the nature of protein modifications.

**For Table of Contents Only:** 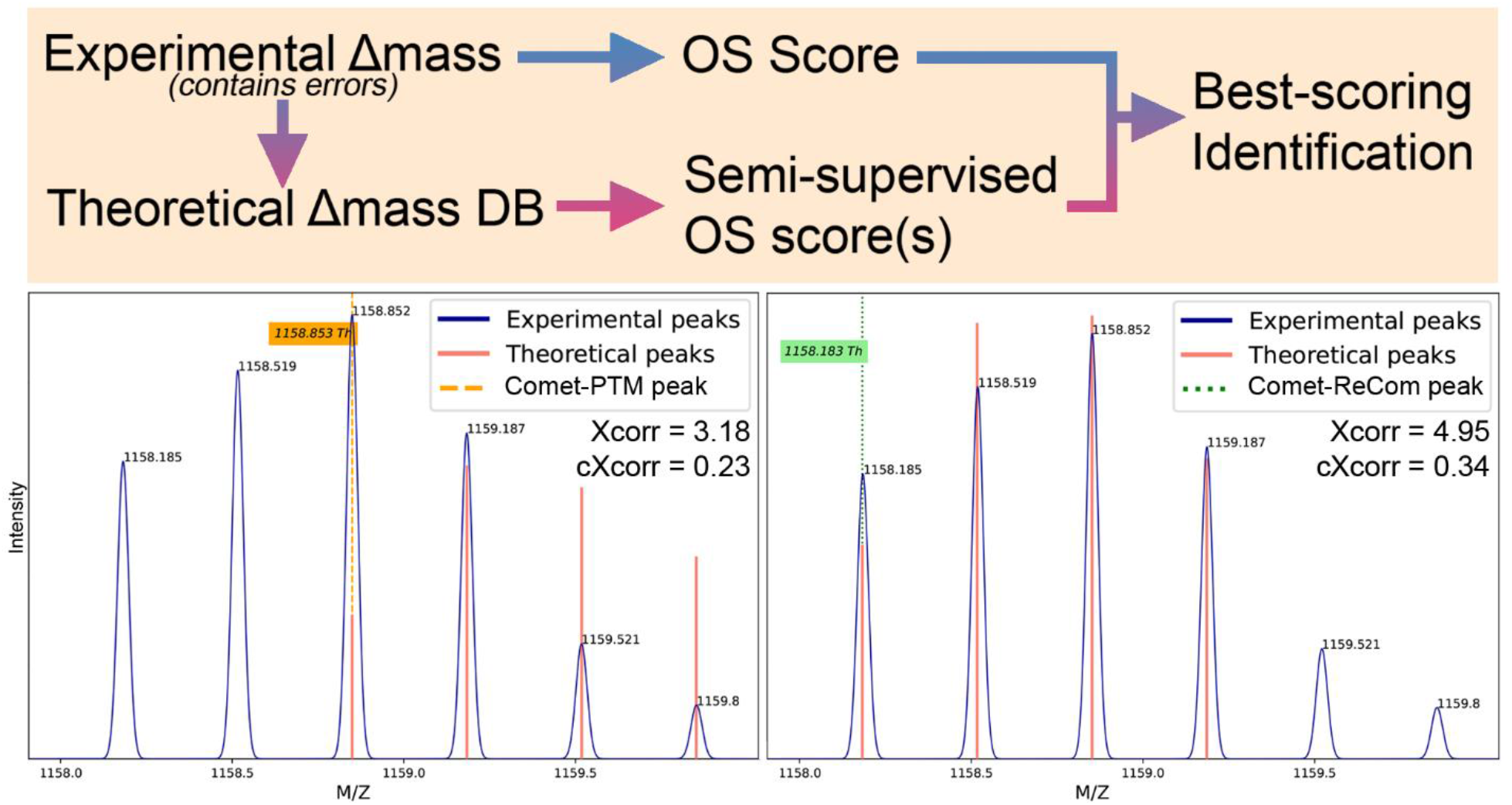

## Introduction

Post-translational modifications (PTMs) substantially increase the functional diversity of proteomes thanks to their ability to regulate protein activity, localization, and interaction. Consequently, they are of paramount importance to regulate signaling pathways. PTM are also involved in many diseases, such as mitochondrial heteroplasmy-associated pathologies, which induce oxidative modifications in the OXPHOS system.(Bagwan, et al., 2018)

Liquid chromatography in tandem with mass spectrometry is one of the most powerful methods for the high-throughput analysis of proteomes. However, a large proportion of spectra remain commonly unidentified, and constitute what has been called the ‘dark matter’ of proteomics.(Skinner and Kelleher, 2015) The reasons for the existence of these unmapped spectra include the presence of unexpected chemical and post-translational modifications, which alter the mass of the precursor peptide that produces the spectrum. For this reason, conventional (or “closed”) database search approaches that use narrow precursor mass tolerances (which usually lie in the ppm range) cannot identify these modified peptides, unless the mass of the modification is included a priori in the search parameters. Hence, closed-search approaches are unable to identify unknown or unexpected modifications.

In order to study the biological significance of the vast diversity of PTMs, several wide-mass range database searching methods have been developed. A commonly used approach is the ‘open-search’ strategy, which uses conventional search engines with ultra-tolerant precursor mass windows (up to ±500 Dalton) to allow matching peptides with modifications. These methods are able to identify modified peptides at an unprecedented scale (Chick, et al., 2015) (Skinner and Kelleher, 2015). The original methods used unmodified fragment ion masses to identify the modified peptides (Chick, et al., 2015; Kong, et al., 2017) and hence were only able to detect half of all potentially detectable modifications.(Bagwan, et al., 2018) Furthermore, the position of the modified residue could not be directly determined.

More recent open-search algorithms such as Comet-PTM (Bagwan, et al., 2018) or MSFragger (Yu, et al., 2020) calculate the mass difference between the observed mass of the peptide ion and the theoretical mass of the unmodified peptide candidate (*Δmass*) to improve identification performance, making it comparable, for specific modifications, to that obtained from closed-searches.(Bagwan, et al., 2018) However, these approaches base their performance on the correct estimation of *Δmass*, which often contain errors. These deviations may be small, often due to measurement errors of the MS machine; or large (i.e. > 0.1 Da), due to incorrect assignation of the precursor mass. Large errors may arise from either the selection of a non-monoisotopic precursor peak from the isotopic envelope of the same precursor peptide, or a peak that belongs to a completely different precursor peptide derived from ion interference of near-isobaric coeluting peptides.

In this work we propose a semi-supervised open search approach called ReCom that minimizes the error in *Δmass* by taking advantage of a priori known information taken from a reference database, such as Unimod or a database provided by the user. We present a proof-of-concept study using an improved version of Comet-PTM, called Comet-ReCom. Our results demonstrate that this semi-supervised approach overcomes the restrictions caused by experimental errors in traditional open search methods and recover a larger amount of useful information from the same dataset, thus improving the performance of PTM identification.

## Methods

### Software Development

We developed a modified version of the Comet-PTM search engine (v2018.01) called Comet-ReCom, which is available at the CNIC-Proteomics GitHub repository (https://github.com/CNIC-Proteomics/ReCom). Comet-ReCom uses a curated list of theoretical mass differences produced by known chemical or posttranslational peptide modifications, selected from the Unimod database. For this study we used a list containing 435 entries obtained by selecting post-translational modifications and chemical derivatives in the -500 to 500 Da range from the Unimod database (Supporting Information, File 1), and reanalyzed only the matches of the two highest-scoring candidates assigned by Comet-PTM to each scan. We included in the reanalysis all theoretical mass differences which were within a range of ±2.3 Dalton. Comet-ReCom used Xcorr to score candidates and to choose the best match.

### MS dataset

The mass spectrometry data used in this study was obtained from the previously published LC-MS/MS analysis of a nucleolar subproteome obtained from peripheral blood lymphocytes from human donors (Martín-Cófreces, et al., 2020) and is accessible from PeptideAtlas at http://peptideatlas.org/PASS/PASS01393. To access files via FTP, use the following credentials: Servername: ftp.peptideatlas.org; Username: PASS01393; Password: YP9856j.

### Data analysis

Database searching of MS data was performed with Comet-PTM or with Comet-ReCom against a human reference proteome concatenated target-decoy database (Supporting Information, File 2) (436237 sequences). Database searching parameters (Supporting Information, File 3) were selected as follows: trypsin digestion with 2 maximum missed cleavage sites, precursor mass tolerance of 2 Da, fragment mass tolerance of 0.03 Da. Cysteine carbamidomethylation (+57.021464 Da) and iTRAQ labeling (+144.102063 Da) at Lysine and peptide N-terminal were considered as fixed modifications.

For Comet-PTM data, recalibration of experimental *Δmass* values was done using a high-quality subset of PTMs as reference (minimum corrected Xcorr 0.15, maximum error 20ppm) as described.(Bagwan, et al., 2018) Peak modelling and peak assignation were also performed as described.(Bagwan, et al., 2018). For PSM identification 1% FDR threshold was used at global, local (1 Da bins) and *Δmass* peak levels. Comet-ReCom reanalysis does not require calibration or peak modelling since the *Δmasses* included in the list are exact theoretical values.

### Comparative analysis of performance of Comet-PTM and Comet-ReCom

To analyze whether the theoretical peptide masses of the candidates proposed by the search engines fitted correctly the experimentally observed isotopic envelope of the precursor, we firstly performed a 7-scan spectrum averaging, including the three previous and the three posterior MS scans. Then, the theoretical isotope intensity distribution of the modified peptide sequence was calculated using a Poisson distribution and the Averagine model (Senko, et al., 1995). The theoretical distribution was fitted to the experimental distribution using apex intensities and by adjusting a scale parameter by iterative least squares. Goodness of fit was estimated by computing the mean-squared error of the residuals and normalizing them by the scale parameter, to obtain a scale-independent MSE estimate. To test whether a fit was correct or not, we selected 100 MS spectra among the highest-scoring PSM that were correctly fitted by visual inspection, and calculated the typical MSE of these validated fits. This typical MSE was used as the null hypothesis for validation of goodness of fit using a χ^2^ test. A fit was considered incorrect when its MSE was significantly higher than the typical MSE. F-test was used to determine whether ReCom provided a significantly better fit than Comet-PTM to the observed isotope intensity distribution.

### Comparative analysis of Comet-ReCom with other search engines

In order to compare the performance of Comet-ReCom against other search engines capable of PTM identification (MSFragger, MetaMorpheus G-PTM-D, and pFind 3), we searched the same subset of data from our nucleolar subproteome dataset against the same target/decoy database indicated above using the recommended default parameters for open search and PTM discovery from each one of these engines. Carbamidomethyl on C and iTRAQ4plex on K and N-term were specified as fixed modifications. The results were filtered at 1% FDR using the same target/decoy strategy for MSFragger and Comet-ReCom, or using the FDR variable calculated by the program in the case of pFind and MetaMorpheus.

## Results and Discussion

### Design of ReCom

The philosophy of the semi-supervised open-search algorithm (ReCom) is schematized in Fig.1. This method relies on a curated list of theoretical mass differences produced by PTM with known biological or chemical relevance, obtained from Unimod or provided by the user. Database searching is firstly performed using conventional open search conditions. For each scan, the n^th^ best candidates specified by the user are selected, and for each of them the *Δmass*, or difference between the observed precursor ion mass and the theoretical mass corresponding to the unmodified peptide candidate, is calculated. ReCom then inspects the curated list and looks for the presence of theoretical *Δmass* values similar to the experimental *Δmass* within a user-specified mass tolerance. The algorithm iteratively calculates a new score using each of the alternative theoretical *Δmass* values, and selects the one that produces the best score. If no theoretical *Δmasses* within the mass tolerance are found in the curated list or none of them improve the original score, then the original *Δmass* is maintained. Hence ReCom identifications are not constrained to the curated list, and novel *Δmasses* which may have biological relevance can still be identified.

**Figure 1.**
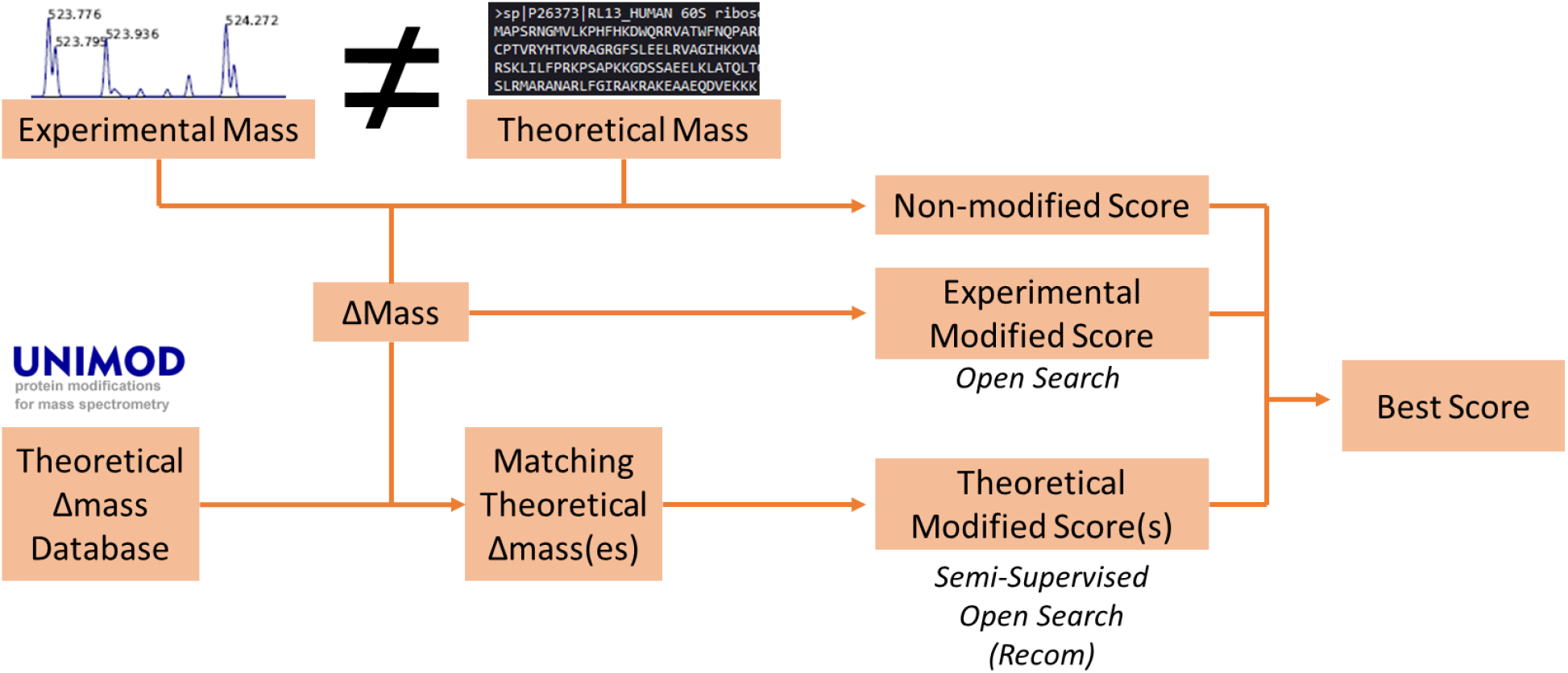
Schematic of the semi-supervised open search strategy employed by ReCom. The curated list of theoretical Δmasses may be taken from an established database or provided by the user. For further details see the text.

### Evaluation of ReCom performance using Comet-PTM

To make a proof-of-concept of the ReCom approach, we modified the source code of the Comet-PTM search engine to re-score peptide candidates following the ReCom algorithm. We used the new engine, Comet-ReCom, with a curated mass list obtained from Unimod as explained in Methods, and reanalyzed a human MS dataset produced in a previous study (Martín-Cófreces, et al., 2020), setting the engine to look for the best two candidates per scan from a concatenated target-decoy sequence database. After 1% FDR filtering we compared the total number of PSMs obtained by Comet-PTM and by Comet-ReCom. Comet-ReCom corrected the *Δmass*, improving the score of a considerable number of target PSMs, with minimal effect on decoy PSMs, allowing to identify 175,986 more PSMs than Comet-PTM, which corresponds to a 68% increase (Fig. 2). In contrast, only 2,838 PSM originally identified by Comet-PTM (less than 1%) were not identified by Comet-ReCom. Hence, ReCom allowed a considerably increase in identification performance in open-search analysis.

**Figure 2.**
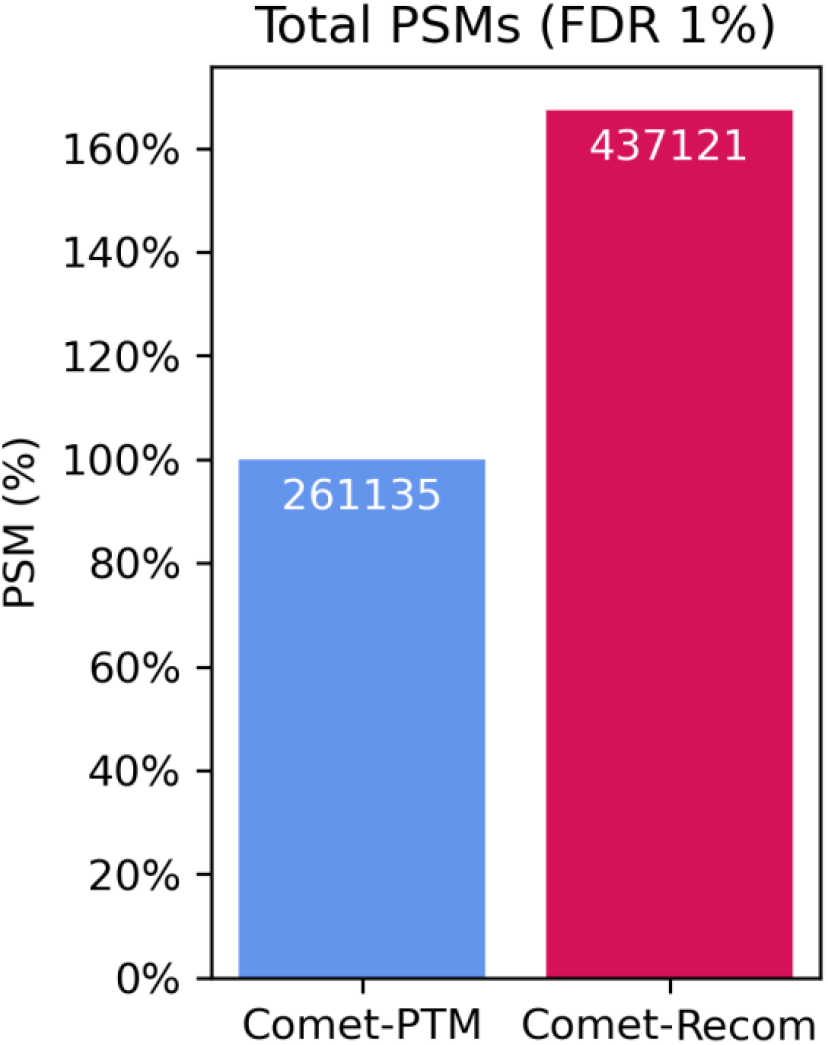
Comparative performance of Comet-PTM and of Comet-ReCom

### Analysis of corrections made by Comet-ReCom

Among the PSM identified by Comet-ReCom, 89,310 of them changed the *Δmass* of Comet-PTM by more than 0.1 Da. In this subset of data, ReCom performed two kinds of *Δmass* corrections: a) isotopic corrections, where ReCom changed the assumed monoisotopic peak to another peak from the isotopic envelope of the same peptide *(Fig. 3A)*, and b) precursor corrections, where ReCom selected a peak that did not belong to the isotopic envelope of the originally matched peptide, pinpointing a different peptide sequence *(Fig. 3B)*.

**Figure 3.**
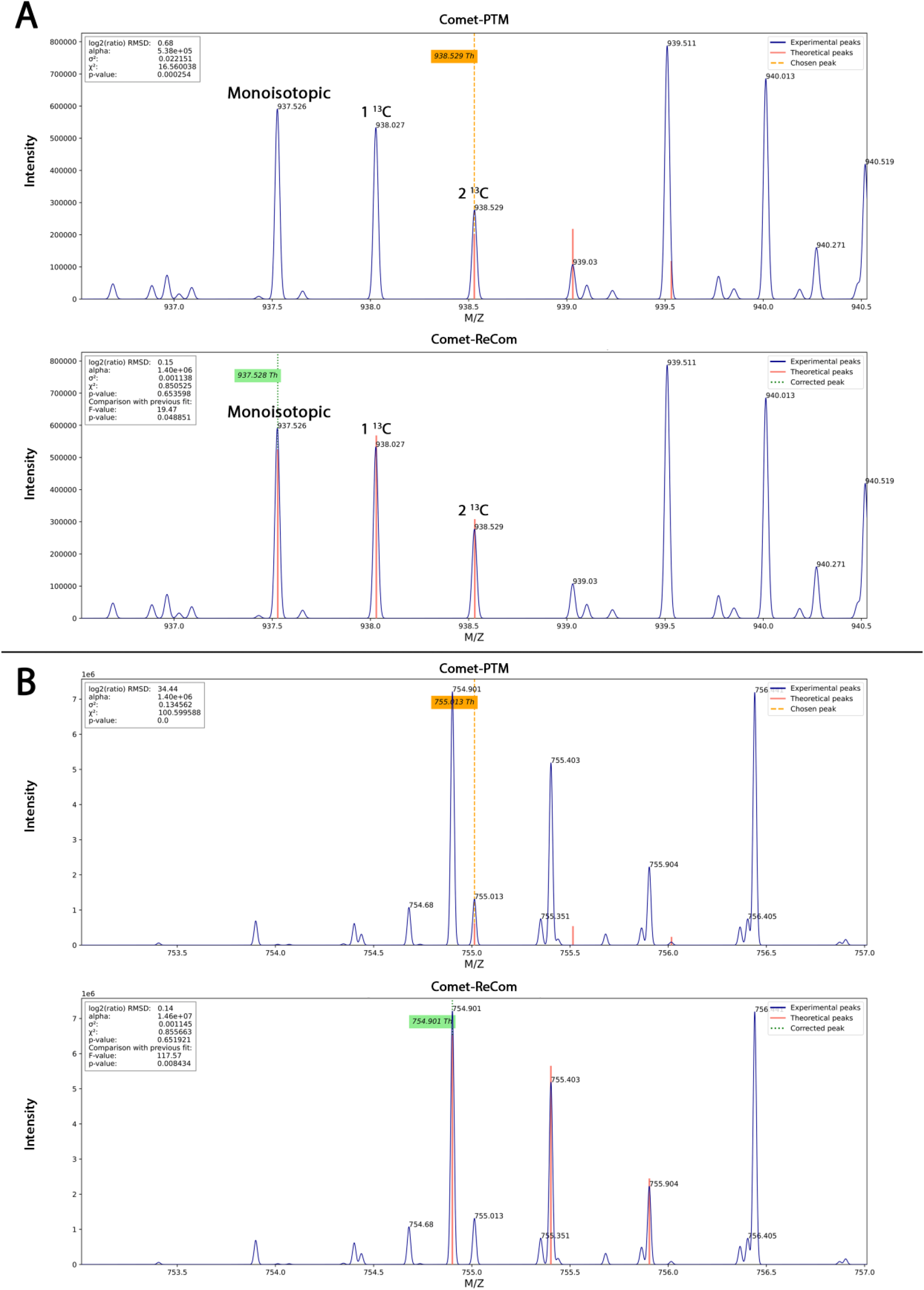
Types of peak corrections performed by ReCom. (A) Example of isotopic correction. The first spectrum indicates the precursor peak chosen by Comet-PTM (yellow), which in fact is the third peak in the isotopic envelope. The second spectrum indicates the precursor peak chosen by ReCom (green) on the sole basis of scoring candidate Δmasses from the theoretical list, which corresponds to the correct monoisotopic peak. (B) Example of precursor correction. In this case, the peak chosen by Comet-PTM is not part of the monoisotopic envelope of the correct precursor, but belongs to a contaminating species, which does not match the experimental distribution. ReCom identifies the correct monoisotopic peak of the precursor.

We observed that corrections were evenly distributed between the isotopic (44.33%) and precursor (55.57%) kinds *(Table 1)*. ReCom performed isotopic corrections jumping over one or two isotopic peaks; corrections over three or more isotopic peaks were not observed because they fell outside of the *Δmass* tolerance range. Precursor corrections of less or more than 1 Da were approximately equally frequent *(Table 1)*.

**Table 1.** Distribution of peak corrections by Comet-ReCom.

|  | PSM | % |
| --- | --- | --- |
| **Corrections** | **89310** | **100.00%** |
| **Isotopic Corrections** | **39681** | **44.43%** |
| └ 1 C13 ($0.99 \leq \text{diff} \leq 1.01$) | 19724 | 22.08% |
| └ 2 C13 ($1.99 \leq \text{diff} \leq 2.01$) | 19957 | 22.35% |
| **Precursor Corrections** | **49629** | **55.57%** |
| └ ($0.1 \leq \text{diff} \leq 0.99$) | 19287 | 21.60% |
| └ ($1.01 \leq \text{diff} \leq 1.99$) | 21930 | 24.55% |
| └ ($2.01 \leq \text{diff}$) | 8412 | 9.42% |

Listed are the corrections where Comet-ReCom made an absolute change in Δmass equal or higher than 0.1 Da.

To further study the nature of isotopic corrections performed by ReCom, we analyzed three well-known modifications that were readily detected in the *Δmass* peak distribution: oxidation, methylation and extra-labeling by the iTRAQ reagent. In the three cases, we found that Comet-ReCom increased by more than 150% the number of PSM whose *Δmass* matched the expected monoisotopic peak (Fig. 5A). In addition, while only approximately half of the Comet-PTM *Δmass* assignations corresponded to the monoisotopic peak, Comet-ReCom assigned the monoisotopic peak in more than 81% of the cases (Fig. 5B). In the case of the iTRAQ peak practically all the PSM were assigned to the monoisotopic peak, probably because the low number of candidate theoretical mass differences for this modification in the curated list within the mass window range. We conclude that ReCom was very efficient in correcting isotopic misassignations.

**Figure 4.**
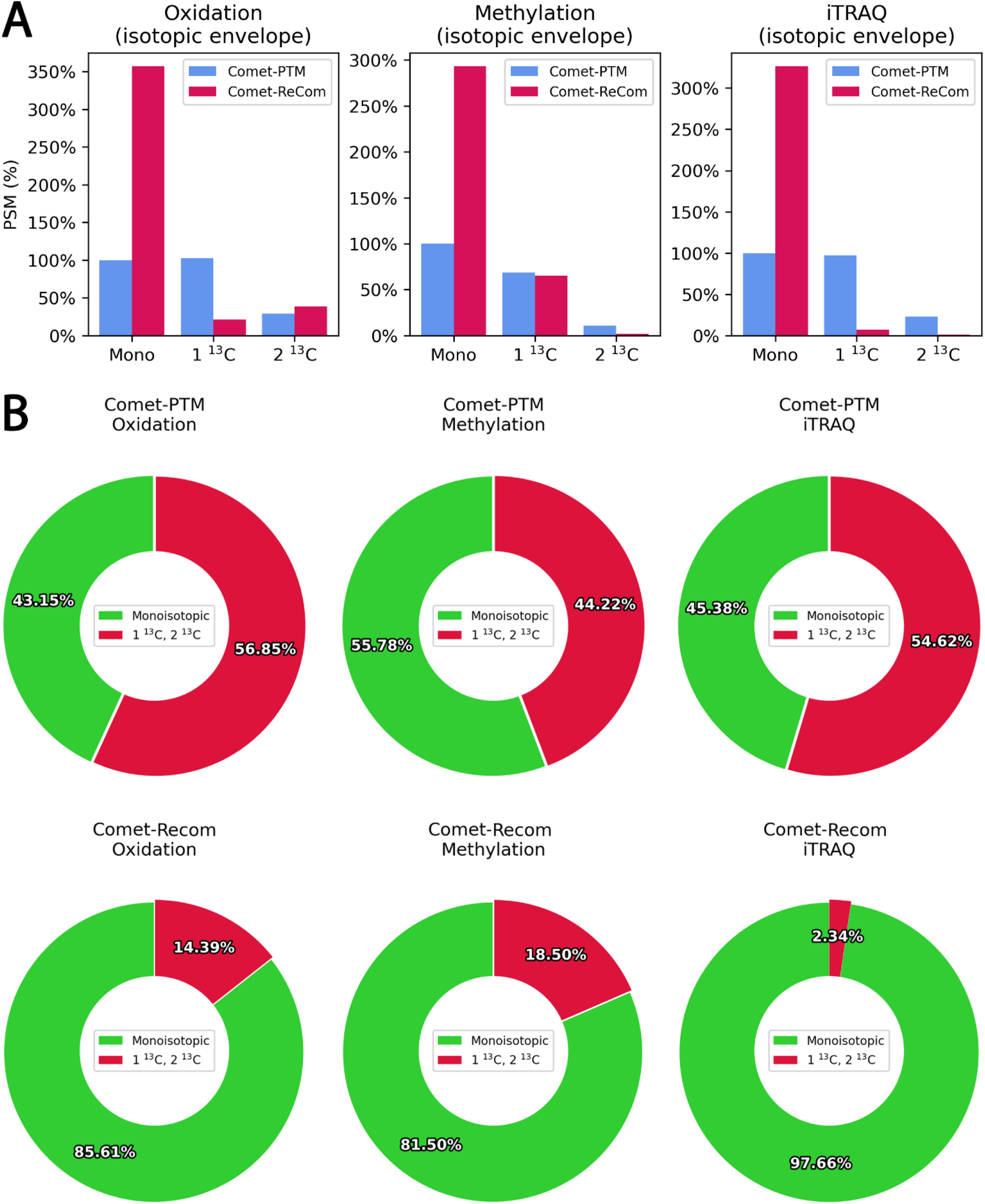
Isotopic corrections performed by Comet-ReCom around three well-known Δmasses. (A) distribution of identifications by Comet-PTM and Comet-ReCom along the three main isotopic peaks. (B) Proportion of assumed monoisotopic assignations for each peak before and after ReCom correction. Δmass assignations were assumed to be monoisotopic when they matched the theoretical mass increment (15.994915, 14.015650 or 144.102063 Da, respectively) and non-monoisotopic when they matched the increments produced by substitution of one or two ^13^C.

**Figure 5.**
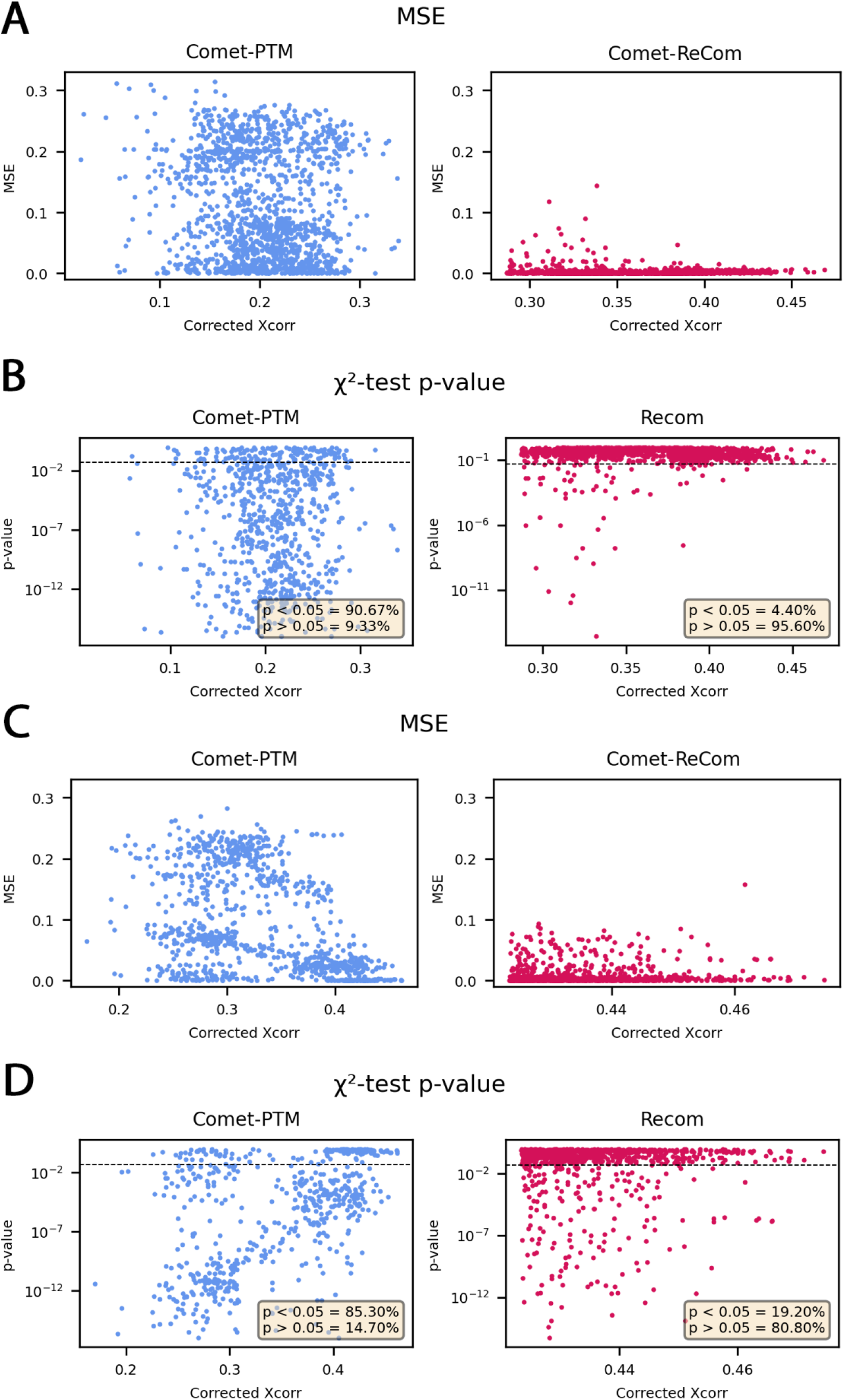
Statistical validation of Comet-ReCom results. The goodness of fit of the theoretical isotope envelope to the observed peaks was measured in terms of the normalized mean squared errors (MSE) (A and C) or by applying a χ^2^-test to determine whether the fitting had a MSE significantly higher than expected (B and D). The analysis was performed on a population containing the 500 highest-scoring PSM from oxidation, methylation and iTRAQ modifications (A and B), or a population containing the 1000 highest-scoring PSM (C and D), whose absolute change in Dmass was higher than 0.1 Da. For further details see the text.

### Statistical validation of Comet-ReCom results

The assignation of the expected monoisotopic peak of known modifications does not fully demonstrate that these assignations are correct. Similarly, precursor corrections can neither be validated on the basis of inspecting *Δmass* values alone. Hence, to further validate the corrections made by Comet-ReCom, we selected the 500 best-scoring PSM from each one of the three common modifications analyzed above where the absolute *Δmass* correction by Comet-ReCom over Comet-PTM was higher than 0.1 Da.

Per each PSM the theoretical isotopic envelope of the precursor peptide ions, generated using the Averagine method and according to the observed *Δmass*, was fitted by least-squares to the peaks present in the MS spectra, and the goodness of fit was analyzed. We observed that the average differences between the observed and fitted intensities (in terms of the mean squared errors of the residuals, or MSE) obtained by Comet-PTM diminished considerably when the results were corrected by Comet-ReCom *(Fig. 5A)*, indicating that Comet-ReCom results fitted better the observed isotopic envelope of precursor ions.

To determine if the assignations of the monoisotopic peak were correct, we applied a χ^2^ test to estimate whether the theoretical isotope distributions according to Comet-PTM *Δmass* or according to Comet-ReCom *Δmass* were significantly different from the observed distributions. In this test, a statistical significance p < 0.05 was considered to indicate a MSE higher than expected and hence an incorrect fit. We found that 91% of Comet-PTM cases were incorrect fits, while only 4% of Comet-ReCom matches passed the test, indicating that the fit was correct in 96% of the cases *(Fig. 5B)*.

Finally, to check that these results were independent on the modification analyzed we selected the 1.000 best-scoring PSM from all the dataset where the absolute *Δmass* correction performed by Comet-ReCom was higher than 0.1 Da and repeated the same analysis. Again, we found that the goodness of fit to the theoretical isotopic envelope improved considerably after ReCom correction (Fig. 5C). Furthermore, Comet-ReCom fits were correct in 81% of the cases, while only 15% of Comet-PTM fits were correct (Fig. 5D). Hence, the majority of ReCom *Δmass* reassignations were in agreement with the observed isotopic envelope.

As further validation, we directly compared the theoretical distributions produced by Comet-PTM with those produced by Comet-ReCom to determine which ones produced a better fit to the observed isotope distribution. We observed that Comet-ReCom produced a significantly better fit (p < 0.05, F-test) than Comet-PTM in 61.8% of the cases in the subset of PSM belonging to the three modifications commented above (Fig. 6A). Comet-ReCom was also significantly better than Comet-PTM in 62.3% of the cases in the dataset with the best 1000 PSM (Fig. 6B).

**Figure 6.**
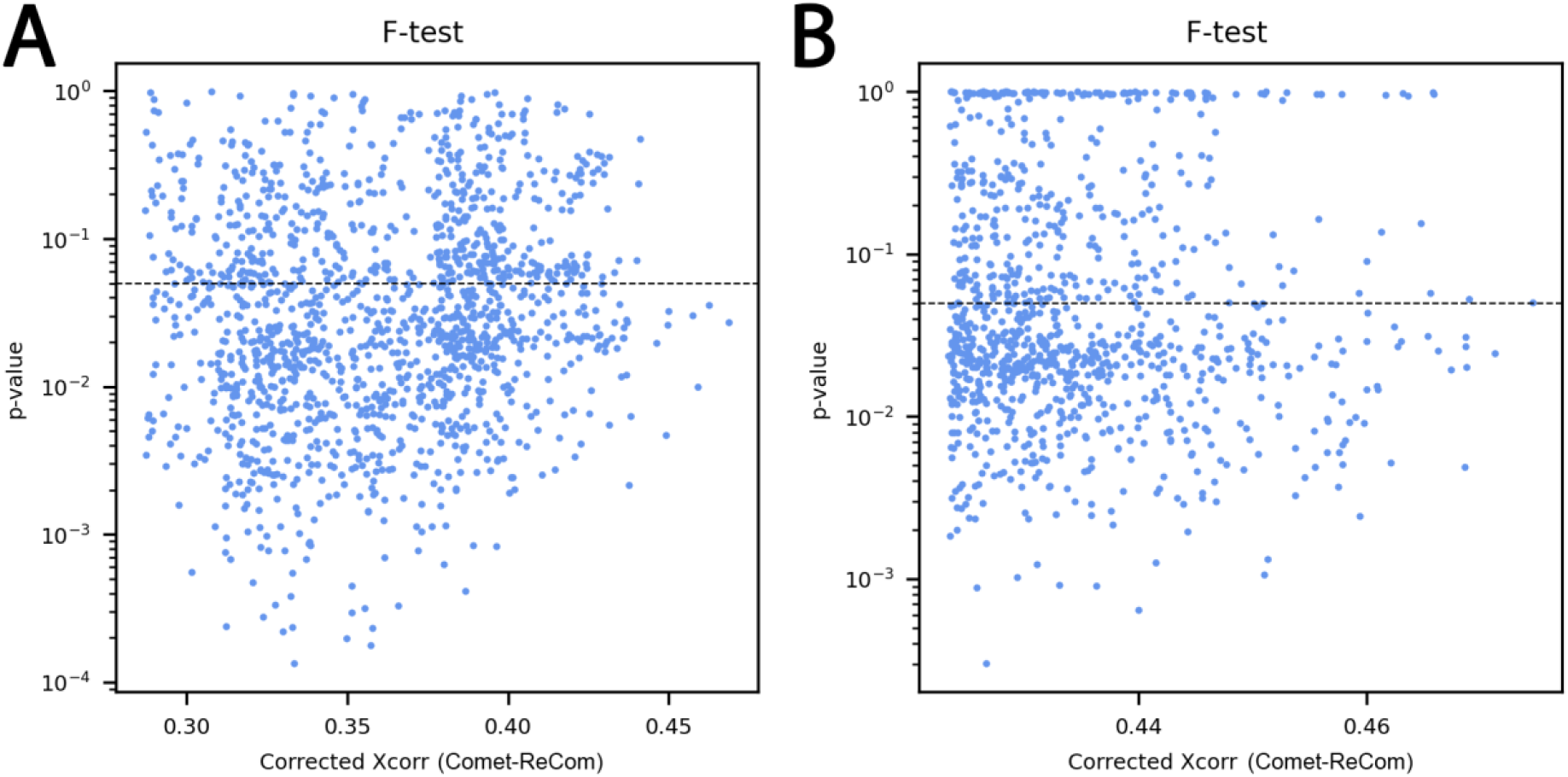
Comparison of the goodness of fit to the observed isotope envelope between Comet-PTM and Comet-ReCom. A and B refer to the PSM populations described in Fig. 5, respectively. When p < 0.05 according to the F-test, the fit by Comet-ReCom was considered to be significantly better than that of Comet-PTM.

The remaining cases where Comet-ReCom did not produce a significantly better fit than Comet-PTM were due to peptide mixtures where the two search engines correctly matched the isotopic envelope of a peptide, but the peptides matched by the searching engines were different (Fig. 7). To determine which of the two search engines provided the most adequate explanation of the observed precursor data, we computed the explained intensity, defined as the sum of intensities of matched peaks in the precursor mass window, as an additional measure of the quality of the match. We found that Comet-ReCom provided a similar or higher explained intensity than Comet-PTM in practically all the cases (Fig. 8A and C). The explained intensity by Comet-ReCom was 1.5-fold or higher than Comet-PTM in 58-88% of cases and 10-fold or higher in 10-28% of cases (Fig. 8B and D), depending on the PSM subset used in the analysis.These results indicate that even in the case of peptide mixtures, ReCom was able to select the peptide candidate which was more likely to be the precursor ion that originated the peaks identified in the MS/MS spectrum.

**Figure 7.**
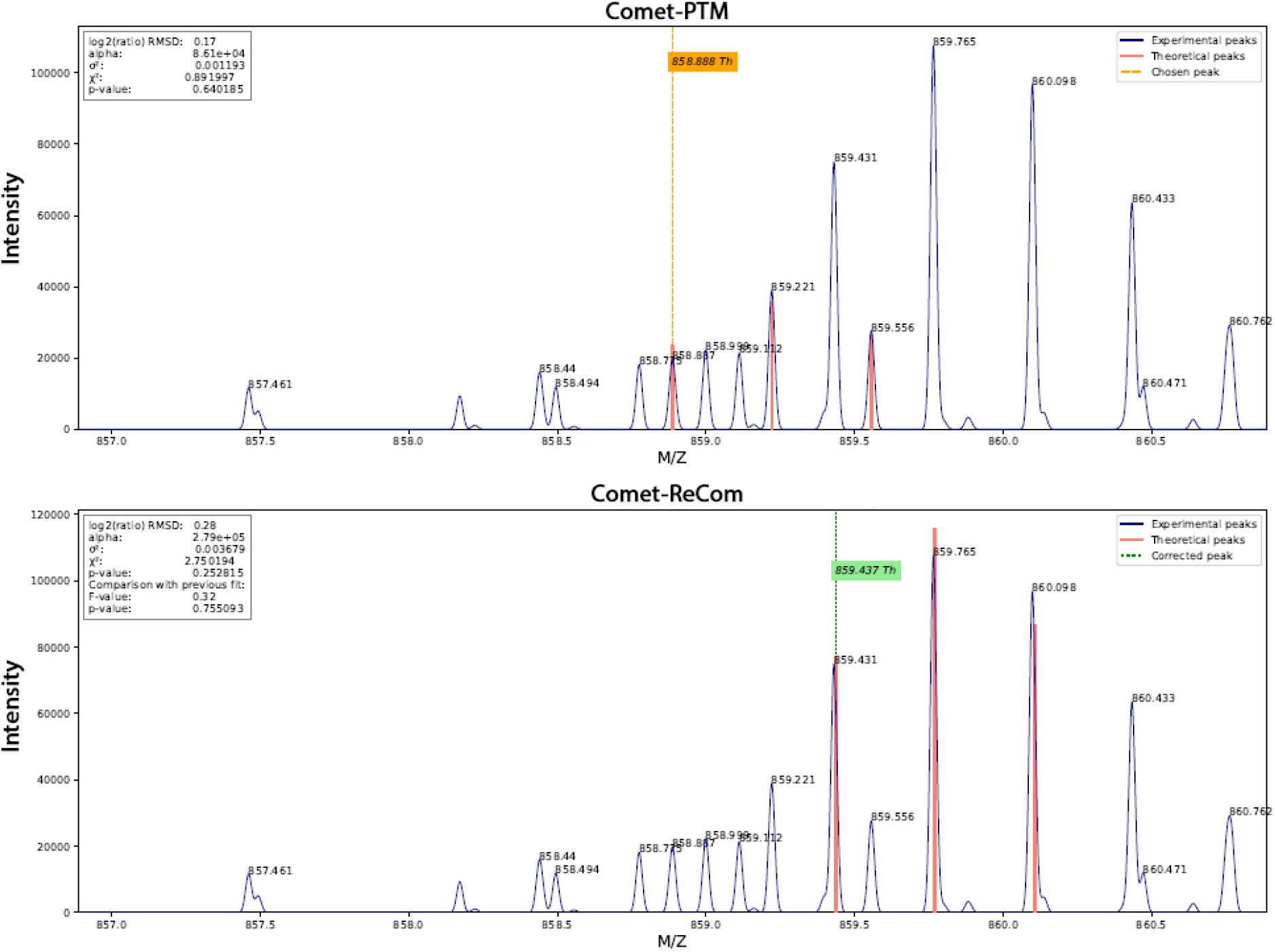
Representative example where both Comet-PTM and ReCom correctly matched the isotopic envelope of a different peptide. Note that Comet-ReCom matched the peptide with higher intensity and hence produced a higher explained intensity (see Fig. 8).

**Figure 8.**
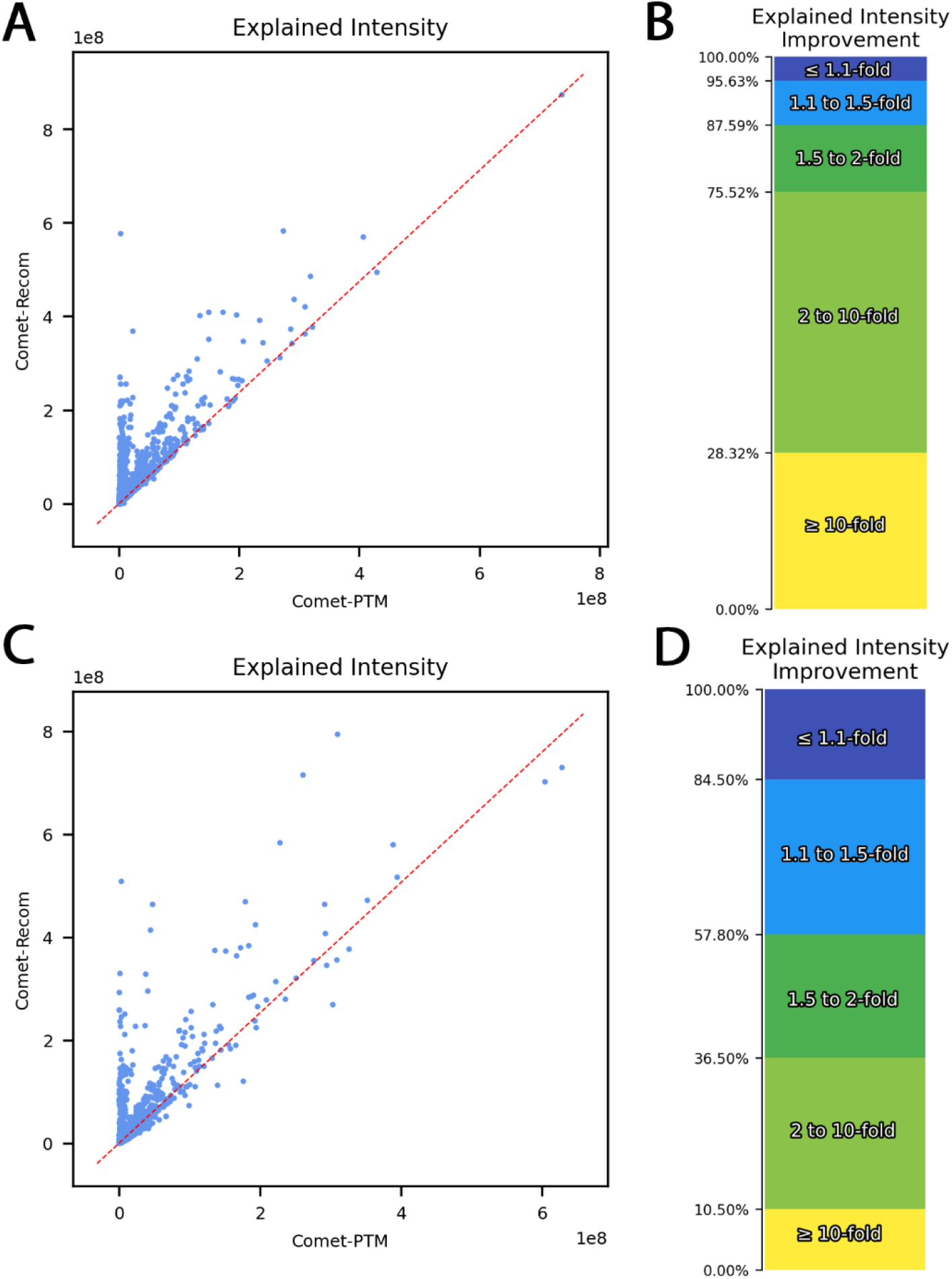
Comparison of explained intensity by Comet-PTM and Comet-ReCom. (A, C) Scatter plot showing that Comet-ReCom consistently achieved an explained intensity similar or greater than Comet-PTM. (B, D) Distribution of explained intensity improvement by Comet-ReCom in comparison with that of Comet-PTM stratified according to the number of PSM. (A, B) and (C, D) refer to the PSM populations described in Fig. 5, respectively.

### Comparison with other search engines for the analysis of PTM

Although the central objective of this paper was to demonstrate the utility of the semi-supervised approach using an open search algorithm (Comet-PTM) whose code was publicly available, we thought it informative to compare its performance with that of other algorithms developed for the analysis of PTM. MSFragger, which is conceptually related to Comet-PTM, performs a localization-aware open search using a conventional scoring algorithm (Yu, et al., 2020). We found that MSFragger managed the problem of identifying the correct isotopic envelope better than Comet-PTM, and the performance of Comet-PTM only became similar to MSFragger when identifications from adjacent isotopic peaks were taken into account (Fig. 9). However, in spite of this improved management, the performance of Comet-ReCom was found to be consistently better than that of MSFragger across different Δmass peaks (Fig. 9). While MSFragger clearly surpasses Comet-ReCom in the practice due to its considerably higher speed, our comparative results suggest that MSFragger could benefit from introducing the semi-supervised philosophy proposed in this work.

**Figure 9.**
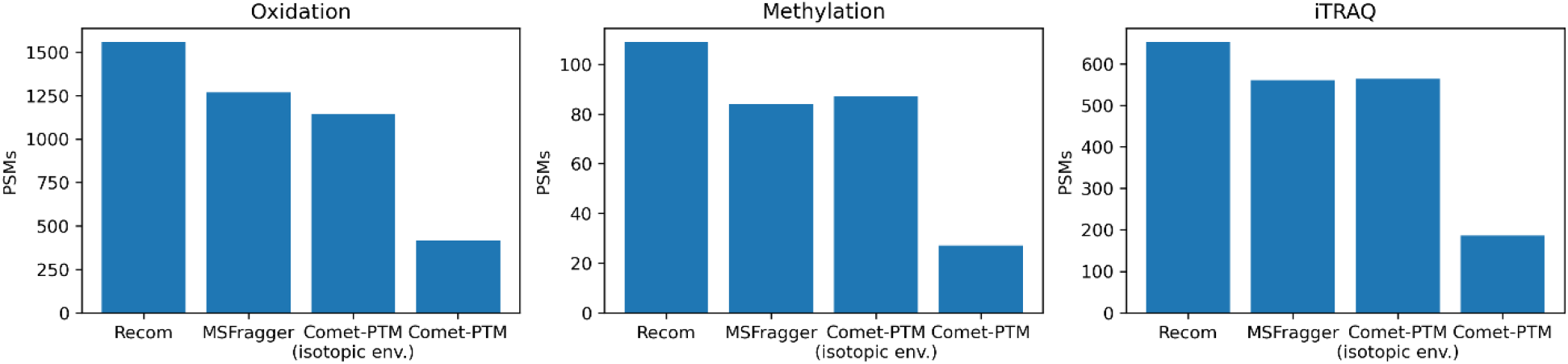
Comparison of identification performance of Comet-PTM, MSFragger and Comet-ReCom. The same dataset was searched against a target/decoy database and results filtered by 1% FDR in all cases. Identification results from three representative examples of Δmass peaks corresponding to the monoisotopic mass increment from each modification are presented. In the case of Comet-PTM we also include, for a more accurate comparison, the performance obtained by adding up identifications in consecutive isotopic peaks (isotopic env.), which would usually be done as a post-processing step.

MetaMorpheus is a proteomics search engine which uses a reference database of theoretical Δmasses to identify modified peptides through its enhanced G-PTM-D (Global Post-translational Modification Discovery) technique (Solntsev, et al., 2018). MetaMorpheus also uses previous information from known modifications; by default this engine considers a set of 66 common biological modifications and artifacts, although different modifications can be selected. In comparison, Comet-ReCom’s default Δmass list contains 435 modifications, and from its design it can also identify Δmass peaks that are not included in this list. We found that while MetaMorpheus was considerably faster than Comet-ReCom and produced similar results for the 40 Δmasses which were identified in its default list, Comet-ReCom was able to further identify 125 theoretical Δmass peaks and additional 10 Δmass peaks of unknown nature (not shown). Hence, MetaMorpheus was more suitable for the fast identification of a limited list of PTM, while Comet-ReCom added the possibility of performing discovery of less-common and novel PTM, at the expense of larger searching times.

pFind 3, with its open search module Open-pFind, is another open search engine which makes use of the Unimod PTM database to identify modified peptides in a semi-supervised manner (Hao Chi, et al., 2018), the main difference being that, unlike Comet-PTM, MSFragger or MetaMorpheus, pFind is based on tandem mass tags to detect potential peptide candidates. When pFind was compared with Comet-ReCom, we observed that pFind consistently produced more identifications in each Δmass peak (Supp Fig. 1A). However, we also found that while there was some overlap, the two search engines identified peptide populations of very different nature. Thus, in the case of oxidations, only 28.5% of them were found by both search engines, while 42.3% of Comet-ReCom oxidations are not found by pFind and 71.4% of pFind identifications were not found by Comet-ReCom (Supp Fig. 1A). A further inspection of these results, however, revealed that Comet-ReCom was often considering pFind candidates among their best-scoring sequences. Thus, if the second oxidation candidate was also considered a match in Comet-ReCom, the overlap between the two engines increased to 66.1% (Supp Fig. 1B). Hence, it was a frequent event than the two engines detected the same candidate in a given MS/MS spectrum, but each one was assigning a higher score to a different candidate. This discrepancy is a consequence of the different approaches used by the two search engines. Oxidized peptides identified uniquely by Comet-ReCom explained a higher proportion of MS/MS peak intensities than the corresponding pFind candidates, as expected for a match and intensity-driven scoring system (Supp Fig. 1B). Similarly, oxidations uniquely identified by pFind did not capture significantly more summed peak intensities from the spectrum than Comet-ReCom candidates, as expected for a tandem mass tag-driven search engine. Some examples of MS/MS spectra where pFind and Comet-ReCom finds different and probably coexisting peptide sequences (Supp. Fig. 2), or which were identified by only one of the two searching engines are presented in Supplementary Material (Supp. Fig. 3). All these results indicate that the two engines produce complementary and mostly different results and suggest that using combined scoring systems could improve identification of modified peptides.

In conclusion, we describe here a new semi-supervised open search approach that addresses one of the limitations of open-search methods that, like MSFragger or Comet-PTM, make use of the precursor mass, obtained from the MS spectrum, to refine the scoring of the MS/MS spectrum. The performance of these search engines is diminished when the precursor mass is not correctly estimated, due for instance to monoisotopic misassignations or to the presence of near isobaric interfering peaks. The ReCom algorithm minimizes this detrimental effect by offering to the search engine an opportunity to match the MS/MS spectrum with the peptide candidate modified using theoretical *Δmasses* similar to the experimentally determined, but which are calculated theoretically from previous knowledge. The number of peptide candidates used for recalculation can be defined by the user, but we show here that by just choosing the two best candidates per scan, increases in peptide identification performance higher than 65% can be obtained. We also show that the increased performance to score the MS/MS spectrum comes in parallel with a significantly better assignation of the monoisotopic peak of the precursor peptide in the MS spectrum when the corrected *Δmass* is used. These results support the validity of the ReCom concept and explains the increased identification performance of the new algorithm. In summary, our data demonstrate that open searches using ultra-tolerant mass windows can be benefited by using a semi-supervised approach that takes advantage from previous knowledge on the nature of protein modifications. We also note that the ReCom concept is fully compatible with other open search algorithms like MSFragger. We foresee that the use of semi-supervised approaches will improve statistical algorithms for the high-throughput analysis of post-translational modifications by mass spectrometry.

## Acknowledgements

This study was supported by competitive grants from the Spanish Ministry of Science, Innovation and Universities (PGC2018-097019-B-I00, PID2021-122348NB-I00, PLEC2022-009235 and PLEC2022-009298), the Instituto de Salud Carlos III (Fondo de Investigación Sanitaria grant PRB3 (PT17/0019/0003-ISCIII-SGEFI / ERDF, ProteoRed), Comunidad de Madrid (IMMUNO-VAR, P2022/BMD-7333) and “la Caixa” Banking Foundation (project codes HR17-00247 and HR22-00253). The CNIC is supported by the Instituto de Salud Carlos III (ISCIII), the Ministerio de Ciencia e Innovación (MCIN) and the Pro CNIC Foundation), and is a Severo Ochoa Center of Excellence (grant CEX2020-001041-S funded by MICIN/AEI/10.13039/501100011033).

