## Supplemental File 3 for "ReCom: A semi-supervised approach to ultra-tolerant database search for improved identification of modified peptides"

# recom_version 2018.01 rev. X
# Comet MS/MS search engine parameters file.
# Everything following the '#' symbol is treated as a comment.

database_name = human_202104_uni-sw-tr.target-decoy.fasta


decoy_search = 0                       # 0=no (default), 1=concatenated search, 2=separate search

num_threads = 44                        # 0=poll CPU to set num threads; else specify num threads directly (max 64)

#
# masses
#
peptide_mass_tolerance = 500
peptide_mass_units = 0                 # 0=amu, 1=mmu, 2=ppm
mass_type_parent = 1                   # 0=average masses, 1=monoisotopic masses
mass_type_fragment = 1                 # 0=average masses, 1=monoisotopic masses
precursor_tolerance_type = 1           # 0=MH+ (default), 1=precursor m/z; only valid for amu/mmu tolerances
isotope_error = 1                      # 0=off, 1=on -1/0/1/2/3 (standard C13 error), 2= -8/-4/0/4/8 (for +4/+8 labeling)

#
# search enzyme
#
search_enzyme_number = 1               # choose from list at end of this params file
num_enzyme_termini = 2                 # valid values are 1 (semi-digested), 2 (fully digested, default), 8 N-term, 9 C-term
allowed_missed_cleavage = 1           # maximum value is 5; for enzyme search

#
# Up to 9 variable modifications are supported
# format:  <mass> <residues> <0=variable/else binary> <max_mods_per_peptide> <term_distance> <n/c-term> <required>
#     e.g. 79.966331 STY 0 3 -1 0 0
#
variable_mod01 = 0.0 M 0 3 -1 0 0

max_variable_mods_in_peptide = 2
require_variable_mod = 0

#
# fragment ions
#
# ion trap ms/ms:  1.0005 tolerance, 0.4 offset (mono masses), theoretical_fragment_ions = 1
# high res ms/ms:    0.02 tolerance, 0.0 offset (mono masses), theoretical_fragment_ions = 0
#
fragment_bin_tol = 0.02                # binning to use on fragment ions
fragment_bin_offset = 0.0              # offset position to start the binning (0.0 to 1.0)
theoretical_fragment_ions = 0          # 0=use flanking peaks, 1=M peak only
use_A_ions = 0
use_B_ions = 1
use_C_ions = 0
use_X_ions = 0
use_Y_ions = 1
use_Z_ions = 0
use_NL_ions = 0                        # 0=no, 1=yes to consider NH3/H2O neutral loss peaks

#
# output
#
output_sqtstream = 0                   # 0=no, 1=yes  write sqt to standard output
output_sqtfile = 0                     # 0=no, 1=yes  write sqt file
output_txtfile = 1                     # 0=no, 1=yes  write tab-delimited txt file
output_pepxmlfile = 0                  # 0=no, 1=yes  write pep.xml file
output_percolatorfile = 0              # 0=no, 1=yes  write Percolator tab-delimited input file
output_outfiles = 0                    # 0=no, 1=yes  write .out files
print_expect_score = 1                 # 0=no, 1=yes to replace Sp with expect in out & sqt
num_output_lines = 5                   # num peptide results to show
show_fragment_ions = 0                 # 0=no, 1=yes for out files only

sample_enzyme_number = 1               # Sample enzyme which is possibly different than the one applied to the search.
                                       # Used to calculate NTT & NMC in pepXML output (default=1 for trypsin).

#
# mzXML parameters
#
scan_range = 0 0                 # start and scan scan range to search; 0 as 1st entry ignores parameter
precursor_charge = 2 5                  # precursor charge range to analyze; does not override any existing charge; 0 as 1st entry ignores parameter
override_charge = 2                # 0=no, 1=override precursor charge states, 2=ignore precursor charges outside precursor_charge range, 3=see online
ms_level = 2                           # MS level to analyze, valid are levels 2 (default) or 3
activation_method = HCD                # activation method; used if activation method set; allowed ALL, CID, ECD, ETD, PQD, HCD, IRMPD

#
# misc parameters
#
digest_mass_range = 400.0 6000.0       # MH+ peptide mass range to analyze
num_results = 500                      # number of search hits to store internally
skip_researching = 1                   # for '.out' file output only, 0=search everything again (default), 1=don't search if .out exists
max_fragment_charge = 3                # set maximum fragment charge state to analyze (allowed max 5)
max_precursor_charge = 5               # set maximum precursor charge state to analyze (allowed max 9)
nucleotide_reading_frame = 0           # 0=proteinDB, 1-6, 7=forward three, 8=reverse three, 9=all six
clip_nterm_methionine = 0              # 0=leave sequences as-is; 1=also consider sequence w/o N-term methionine
spectrum_batch_size = 5000                # max. # of spectra to search at a time; 0 to search the entire scan range in one loop
decoy_prefix = DECOY_                  # decoy entries are denoted by this string which is pre-pended to each protein accession
output_suffix =                        # add a suffix to output base names i.e. suffix "-C" generates base-C.pep.xml from base.mzXML input
mass_offsets =                         # one or more mass offsets to search (values substracted from deconvoluted precursor mass)

#
# spectral processing
#
minimum_peaks = 10                     # required minimum number of peaks in spectrum to search (default 10)
minimum_intensity = 0                  # minimum intensity value to read in
remove_precursor_peak = 0              # 0=no, 1=yes, 2=all charge reduced precursor peaks (for ETD)
remove_precursor_tolerance = 1.5       # +- Da tolerance for precursor removal
clear_mz_range = 0.0 0.0               # for iTRAQ/TMT type data; will clear out all peaks in the specified m/z range


#
# CNIC / comet-iq / comet-PTM specific
#
# do deltaX search for: (-delta_tolerance_outer < deltaMass < -delta_tolerance_inner) OR
#                       (+delta_tolerance_inner < deltaMass < +delta_tolerance_outer)
#
use_delta_xcorr = 1                    # 0=no (default), 1=yes
delta_outer_tolerance = 230            # ignored if use_delta_xcorr 0, default 320 
delta_inner_tolerance = 0              # ignored if use_delta_xcorr 0, default 0.8                        
use_delta_back_jumps = 0               # 0=no (default), 1=yes
use_delta_forward_jumps = 0            # 0=no (default), 1=yes
dont_calc_pseudo_non_mod = 1
use_delta_closest = 1                  # 0=no (default), 1=yes
deltamass_peaks = Theoretical_DeltaMass_List.txt  # if use_delta_closest = 1, path must be set here to peaks file
deltamass_peaks_tolerance = 2.3        # +- Da tolerance for matching experimental masses to the list, default 1
cand_number = 2                        # number of candidates to rescore for each identification, default 1
use_xcorr_corr = 1

#
# additional modifications
#

add_Cterm_peptide = 0.0
add_Nterm_peptide = 144.102063
add_Cterm_protein = 0.0
add_Nterm_protein = 0.0

add_G_glycine = 0.0000                 # added to G - avg.  57.0513, mono.  57.02146
add_A_alanine = 0.0000                 # added to A - avg.  71.0779, mono.  71.03711
add_S_serine = 0.0000                  # added to S - avg.  87.0773, mono.  87.03203
add_P_proline = 0.0000                 # added to P - avg.  97.1152, mono.  97.05276
add_V_valine = 0.0000                  # added to V - avg.  99.1311, mono.  99.06841
add_T_threonine = 0.0000               # added to T - avg. 101.1038, mono. 101.04768
add_C_cysteine = 57.021464             # added to C - avg. 103.1429, mono. 103.00918
add_L_leucine = 0.0000                 # added to L - avg. 113.1576, mono. 113.08406
add_I_isoleucine = 0.0000              # added to I - avg. 113.1576, mono. 113.08406
add_N_asparagine = 0.0000              # added to N - avg. 114.1026, mono. 114.04293
add_D_aspartic_acid = 0.0000           # added to D - avg. 115.0874, mono. 115.02694
add_Q_glutamine = 0.0000               # added to Q - avg. 128.1292, mono. 128.05858
add_K_lysine = 144.102063              # added to K - avg. 128.1723, mono. 128.09496
add_E_glutamic_acid = 0.0000           # added to E - avg. 129.1140, mono. 129.04259
add_M_methionine = 0.0000              # added to M - avg. 131.1961, mono. 131.04048
add_O_ornithine = 0.0000               # added to O - avg. 132.1610, mono  132.08988
add_H_histidine = 0.0000               # added to H - avg. 137.1393, mono. 137.05891
add_F_phenylalanine = 0.0000           # added to F - avg. 147.1739, mono. 147.06841
add_R_arginine = 0.0000                # added to R - avg. 156.1857, mono. 156.10111
add_Y_tyrosine = 0.0000                # added to Y - avg. 163.0633, mono. 163.06333
add_W_tryptophan = 0.0000              # added to W - avg. 186.0793, mono. 186.07931
add_B_user_amino_acid = 0.0000         # added to B - avg.   0.0000, mono.   0.00000
add_J_user_amino_acid = 0.0000         # added to J - avg.   0.0000, mono.   0.00000
add_X_user_amino_acid = 0.0000         # added to X - avg.   0.0000, mono.   0.00000
add_Z_user_amino_acid = 0.0000         # added to Z - avg.   0.0000, mono.   0.00000

#
# COMET_ENZYME_INFO _must_ be at the end of this parameters file
#
[COMET_ENZYME_INFO]
0.  No_enzyme              0      -           -
1.  Trypsin                1      KR          P
2.  Trypsin/P              1      KR          -
3.  Lys_C                  1      K           P
4.  Lys_N                  0      K           -
5.  Arg_C                  1      R           P
6.  Asp_N                  0      D           -
7.  CNBr                   1      M           -
8.  Glu_C                  1      DE          P
9.  PepsinA                1      FL          P
10. Chymotrypsin           1      FWYL        P
