## Supplemental Figure 1 for "ReCom: A semi-supervised approach to ultra-tolerant database search for improved identification of modified peptides"

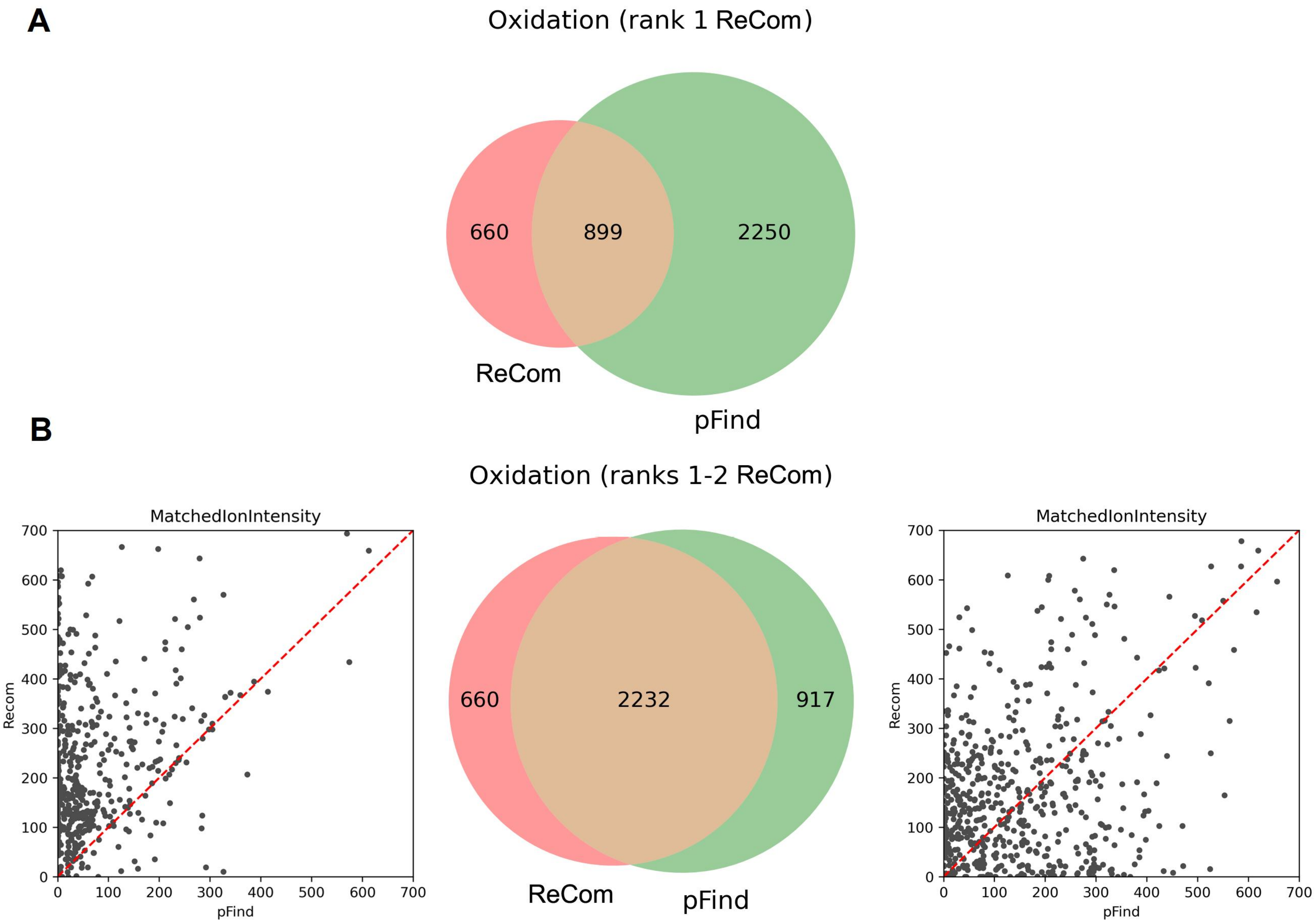


Supplemental Figure 1. Comparison of PSMs containing oxidation identified by Comet-ReCom and pFind, considering only the best candidate found by Comet-ReCom (A) or the best two candidates (B). The sum of the intensities of matched ions was calculated for oxidations identified only by Comet-ReCom vs pFind's alternative identifications (B, left) and for oxidations identified only by pFind vs Comet-ReCom's alternative identifications (B, right)
