## Supplemental Figure 2 for "ReCom: A semi-supervised approach to ultra-tolerant database search for improved identification of modified peptides"

**A**

| | SCAN INFO |
|---|---|
| Raw | JAL_NOa2_iTR_Fr1 |
| Scan | 70721 |
| Charge | 3 |
| RT | 0 |
| DeltaM | 0.016532 |
| Label | iTRAQ4plex |
| MH | 2480.173807 |
| E–score | 0.73865 |

### pFind Candidate

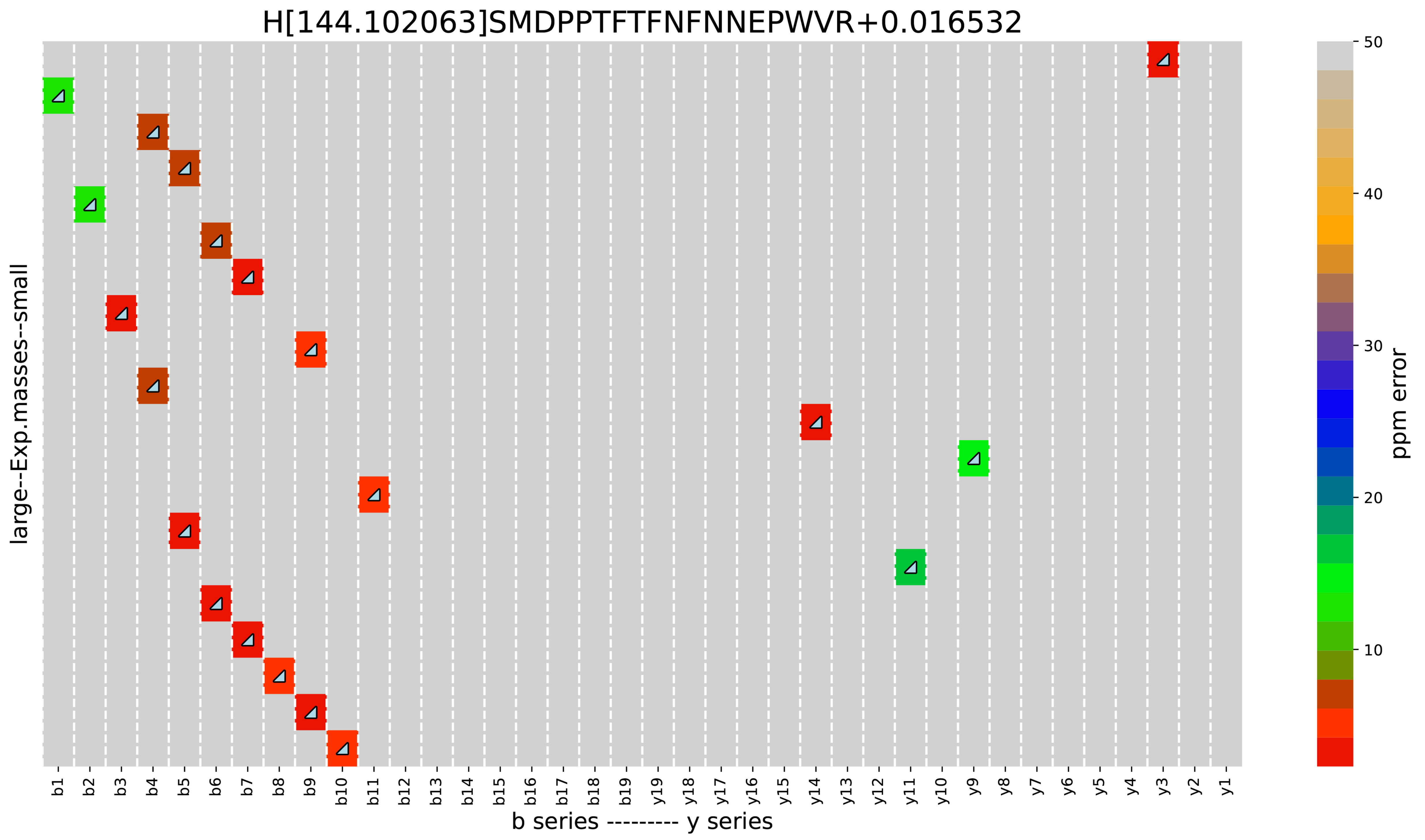


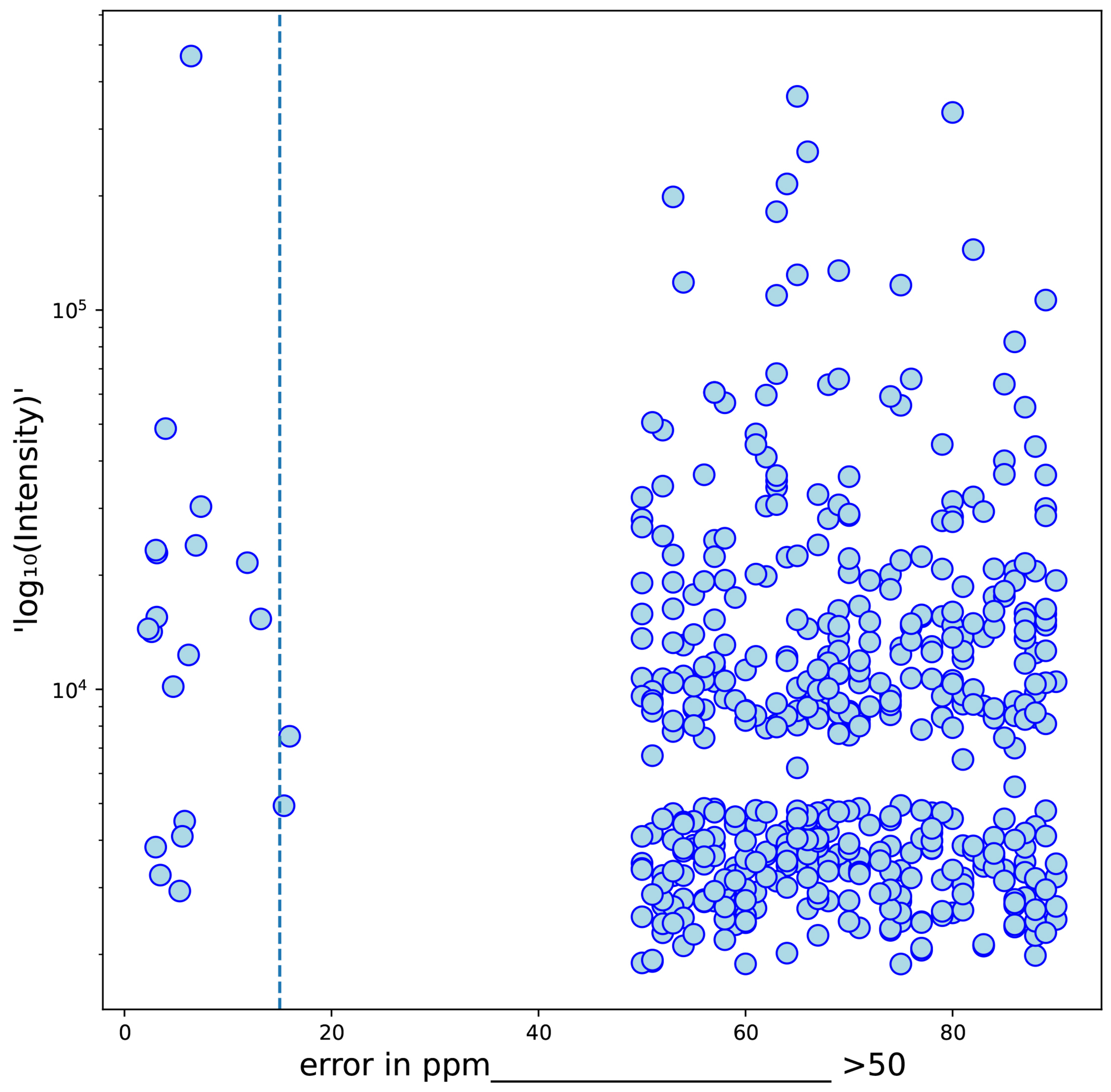


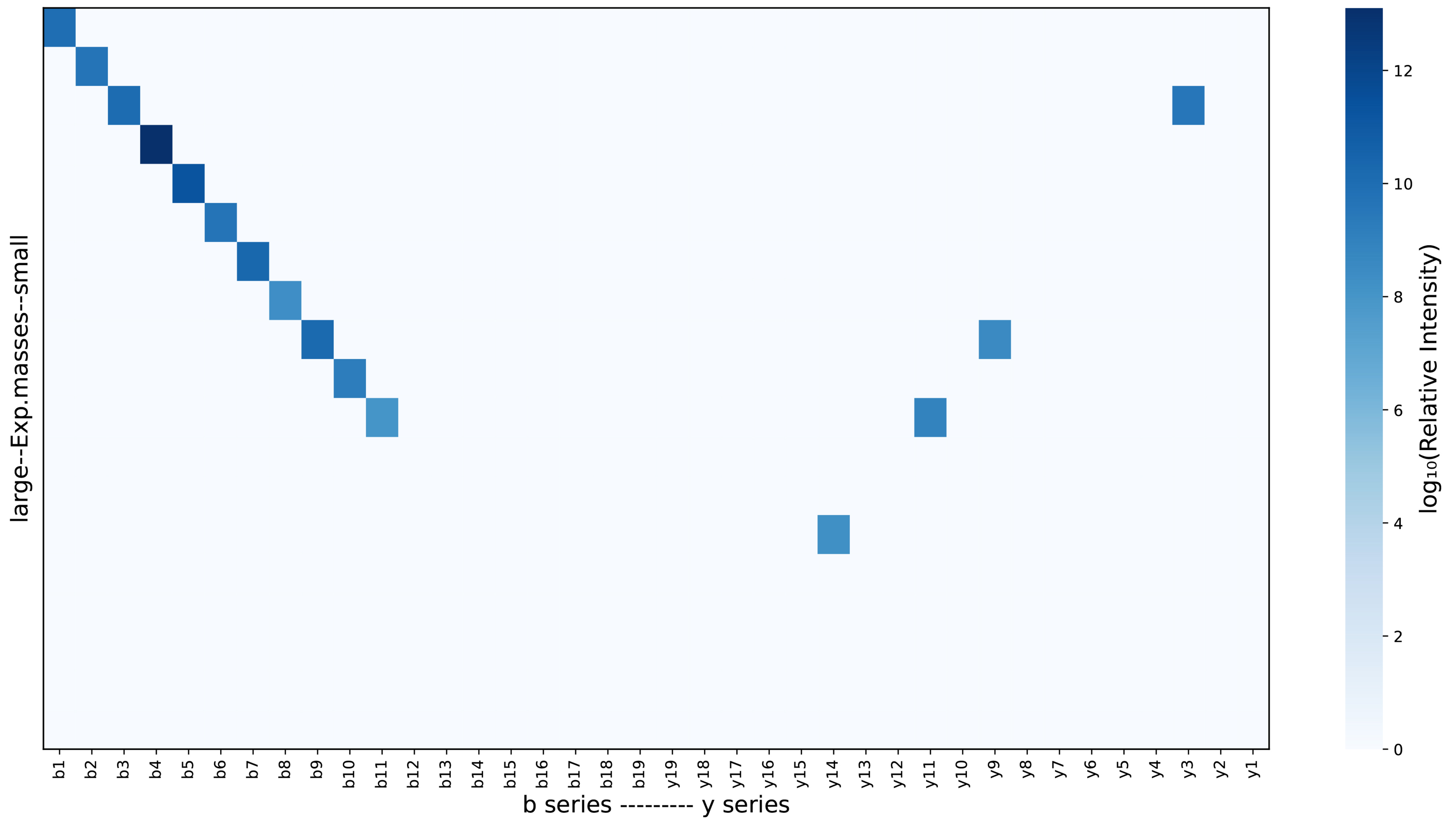


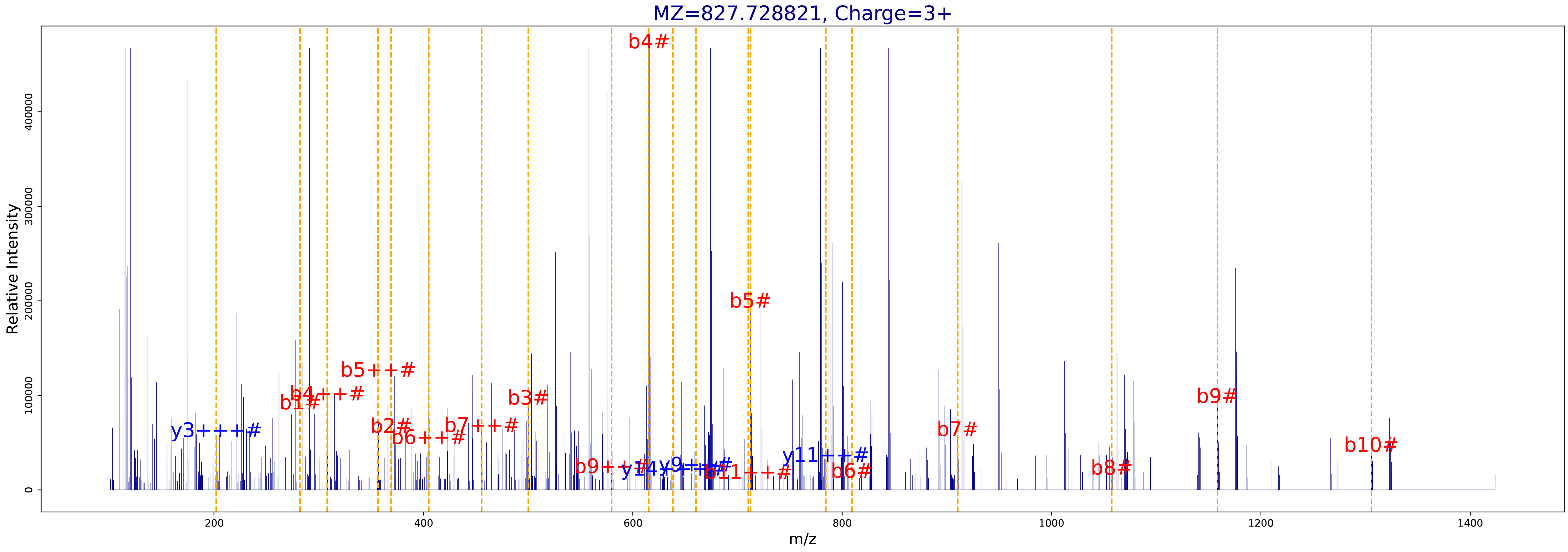

**B**

| | SCAN INFO |
|---|---|
| Raw | JAL_NOa2_iTR_Fr1 |
| Scan | 70721 |
| Charge | 3 |
| RT | 223.79 |
| DeltaM | 15.995138 |
| Label | iTRAQ4plex, iTRAQ4plex |
| MH | 2481.171909 |
| E-score | 1.546191 |

### Recom Candidate

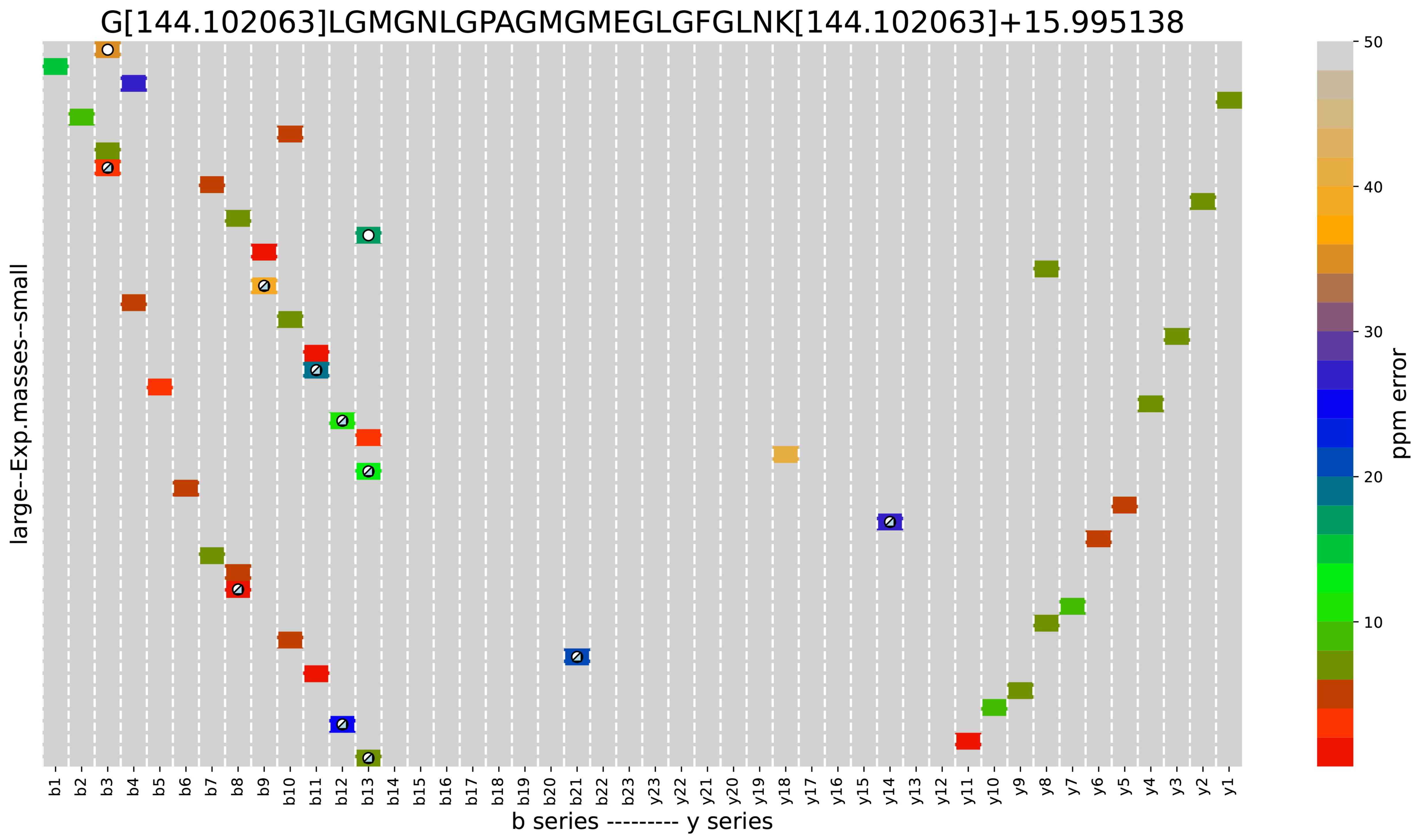


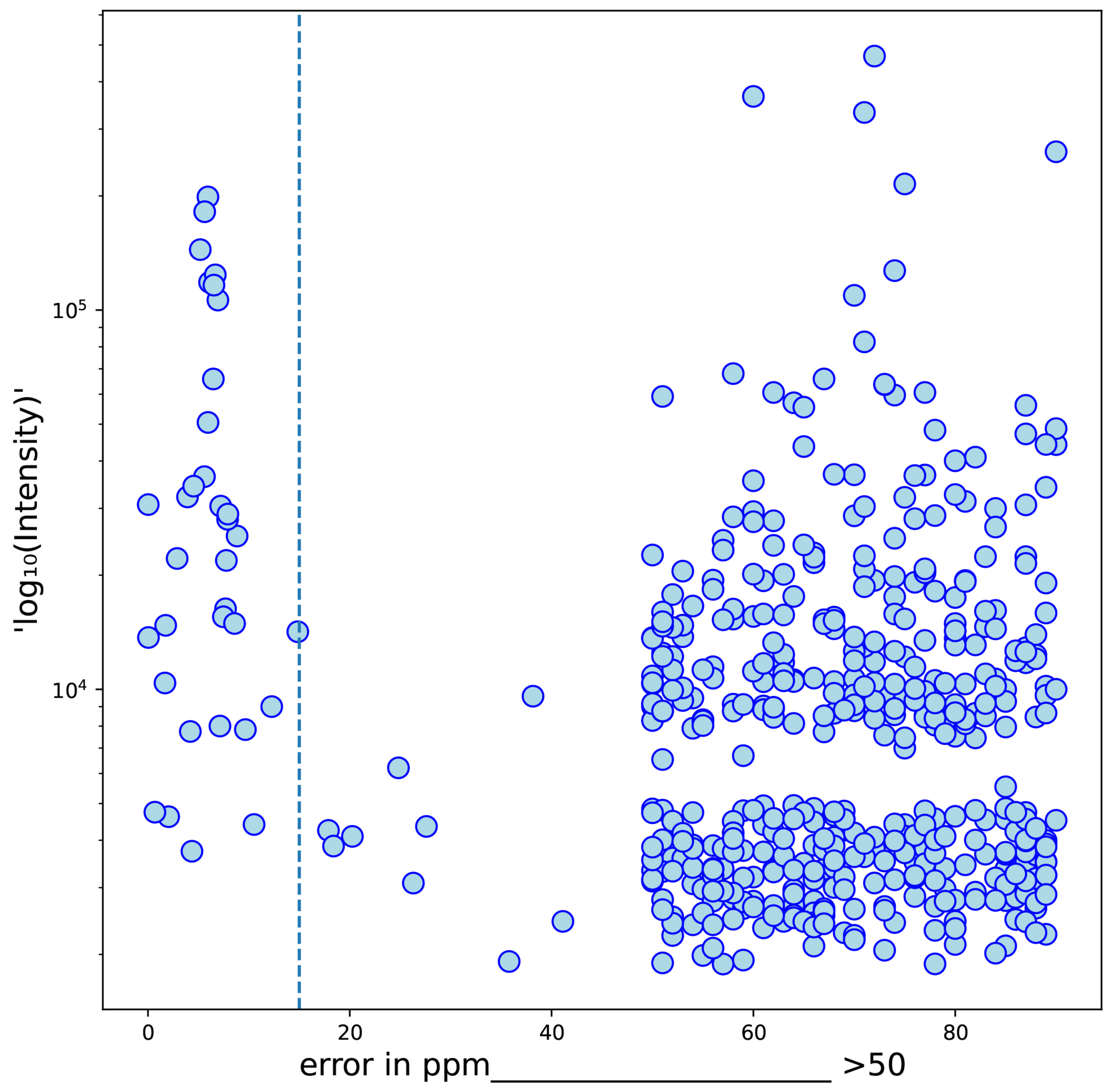


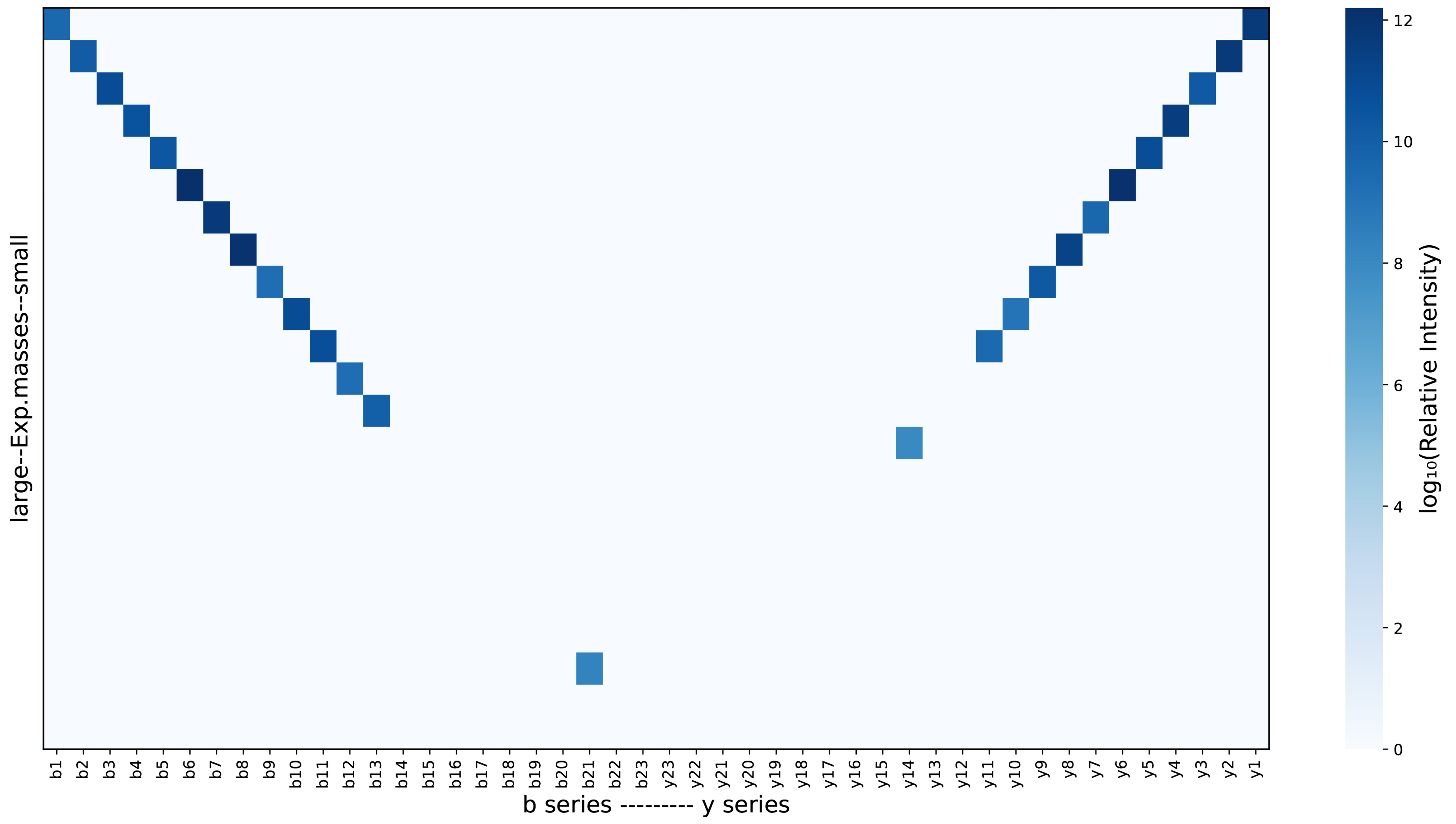


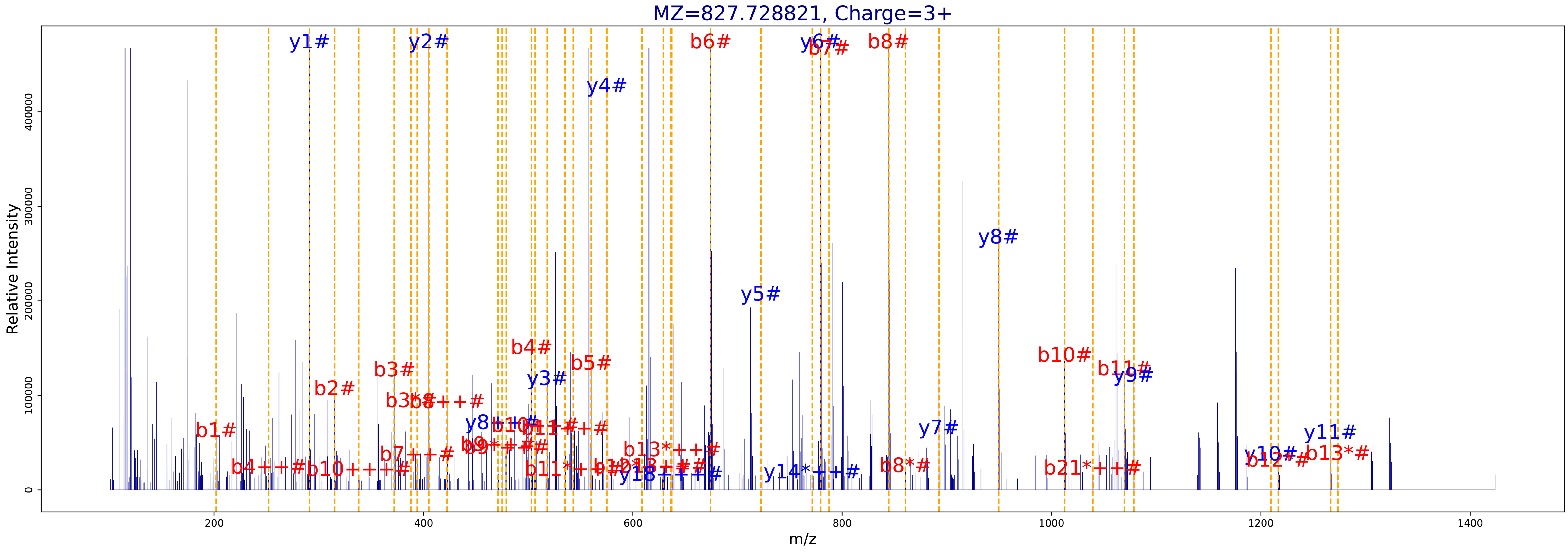

Supplemental Figure 2. Analysis of a different peptide identification by pFind (A) and Comet-ReCom (B) on the same spectrum using Vseq. Vseq (Calvo, et al., 2016) plots show matched fragments (all charges) along the b and y series, colored by their error in ppm and marked with a triangle if they contain a fixed modification and a circle if they contain a ΔMass (upper right panel), a plot of log-intensity against ppm error of fragments (middle left), sequence coverage along the b and y series (middle right), and the assignation of fragments to ion peaks in the spectrum (bottom panel). The two identifications are likely coexisting peptide sequences, a non-modified peptide found by pFind and a peptide containing oxidation on M found by Comet-ReCom. Comet-ReCom has chosen an identification with a longer sequence than pFind, and a greater number of matched fragments below the ppm threshold. Vseq plots show the sequence has high coverage along the b series in the case of pFind, and both the b and y series in the case of Comet-ReCom.
