## Supplemental Figure 3 for "ReCom: A semi-supervised approach to ultra-tolerant database search for improved identification of modified peptides"

**A**

| | SCAN INFO |
|---|---|
| Raw | JAL_NOa2_iTR_Fr1 |
| Scan | 65456 |
| Charge | 4 |
| RT | 205.07 |
| DeltaM | 15.995138 |
| Label | iTRAQ4plex |
| MH | 3343.725973 |
| E-score | 2.225149 |

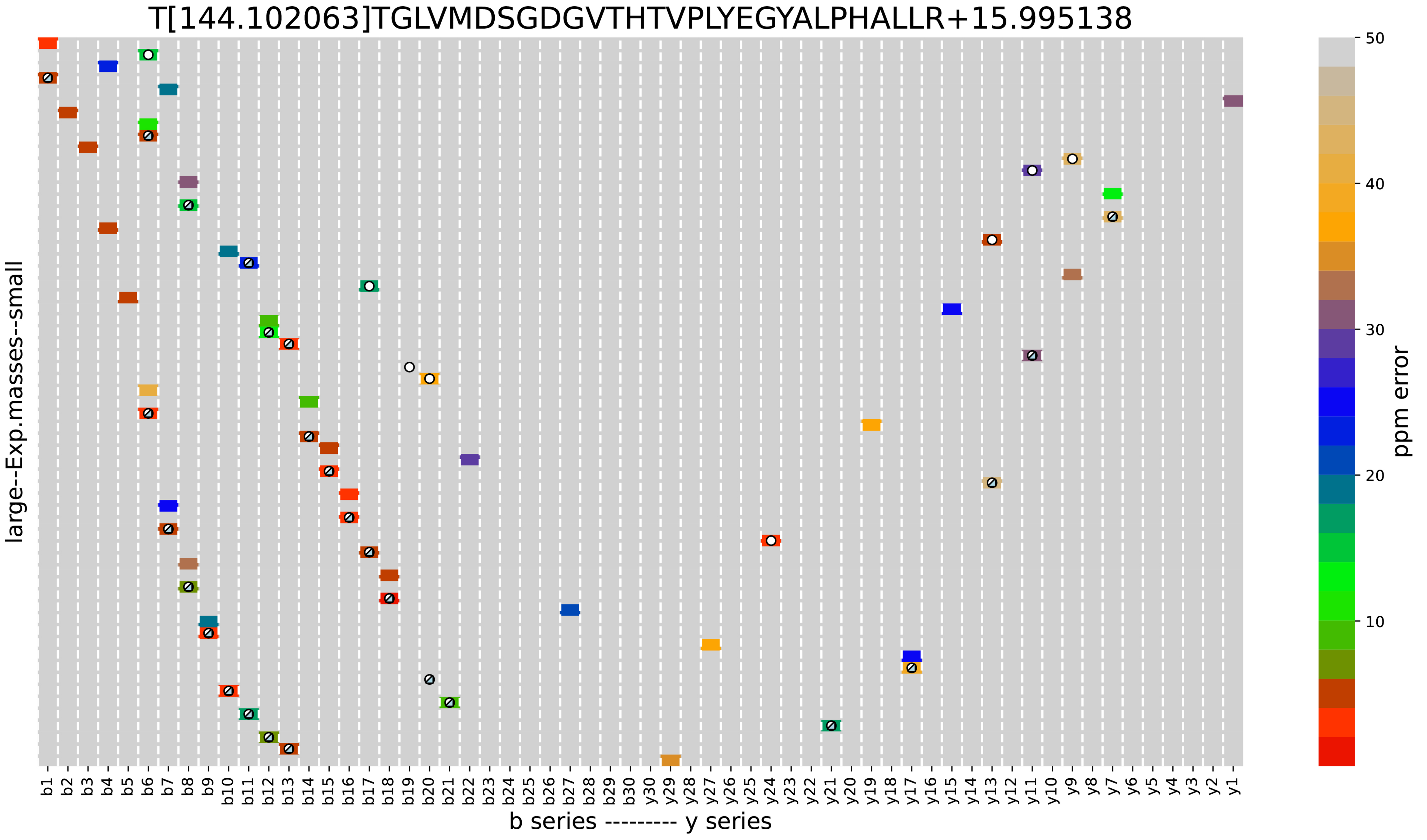


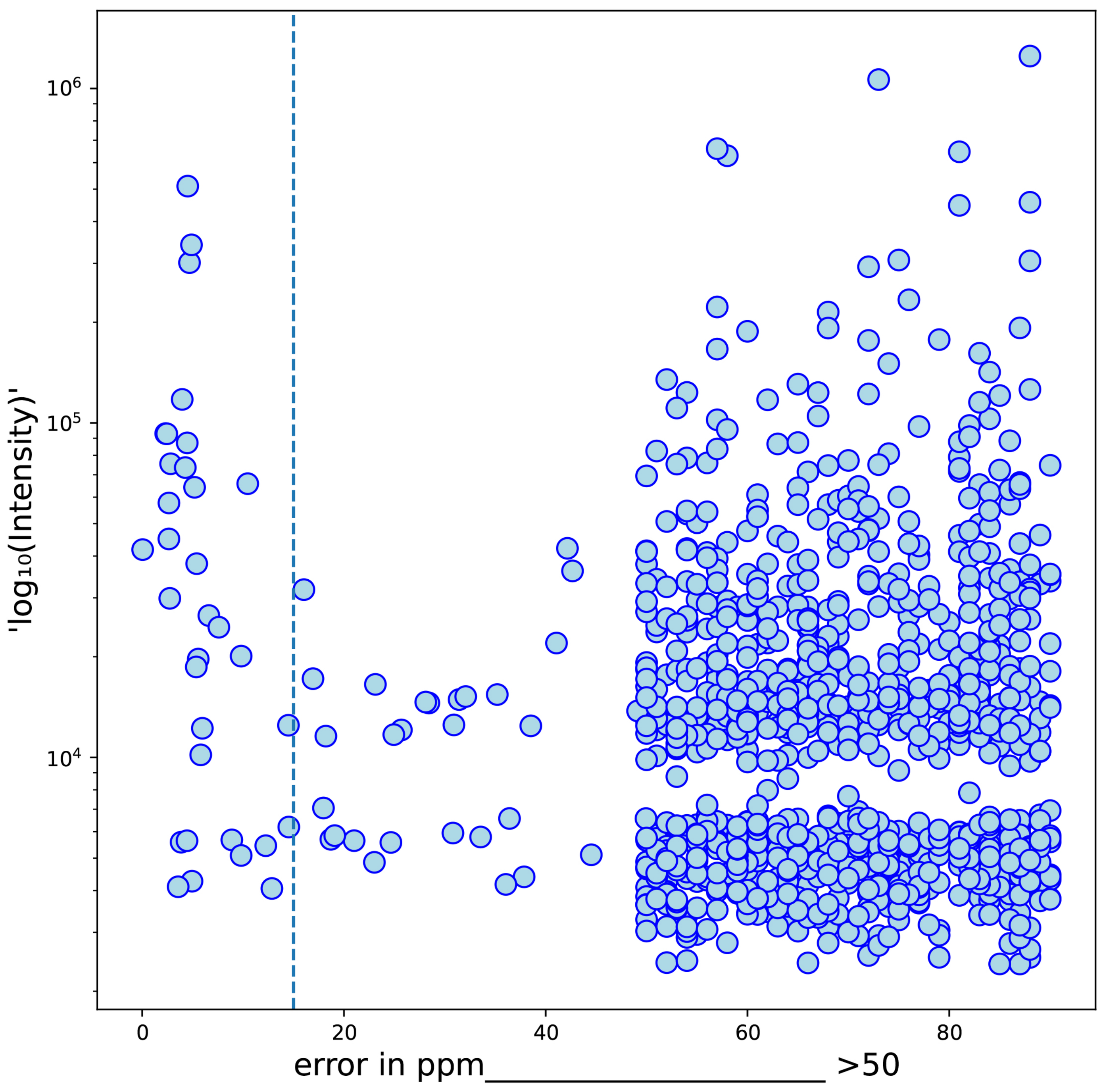


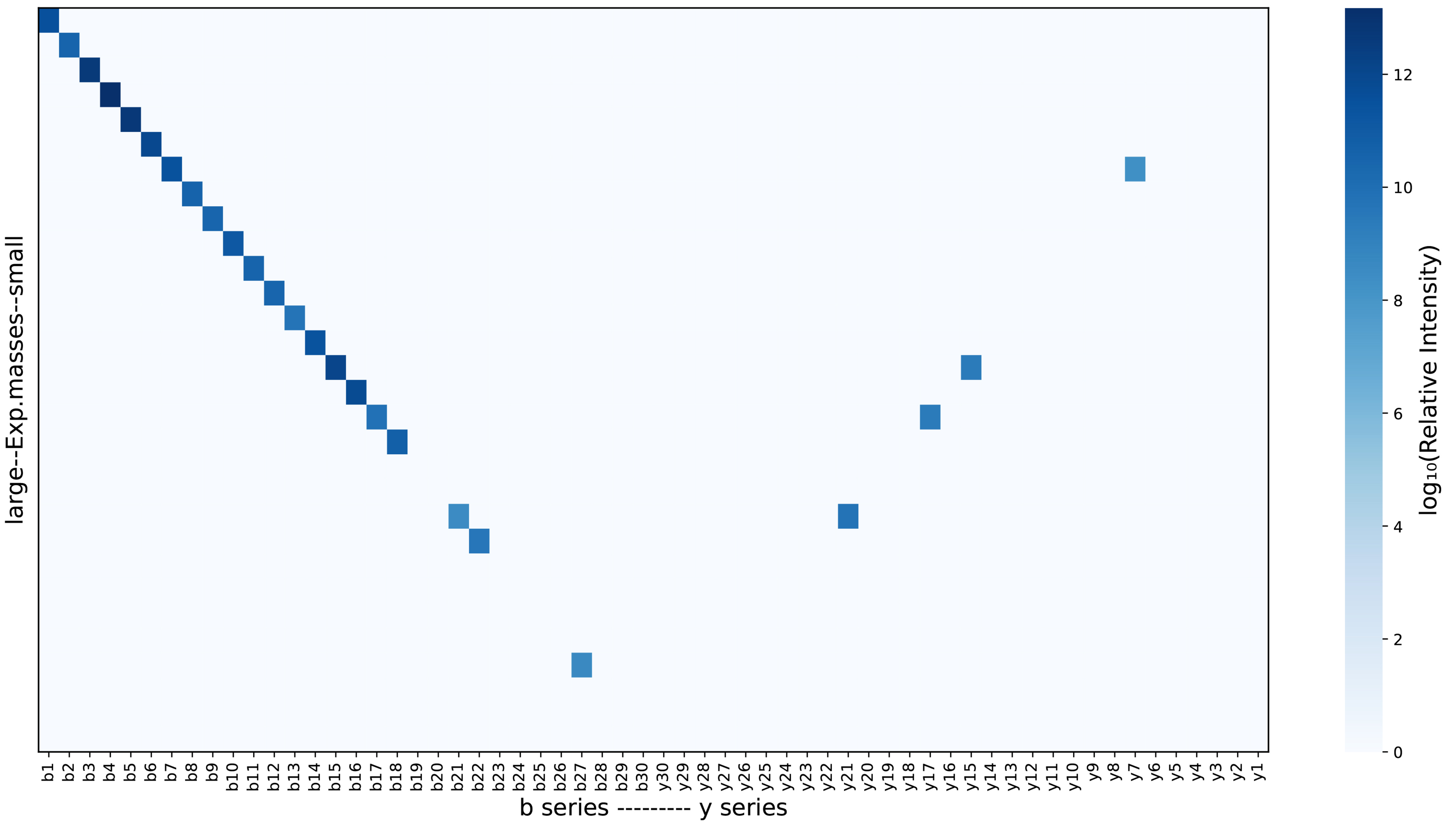


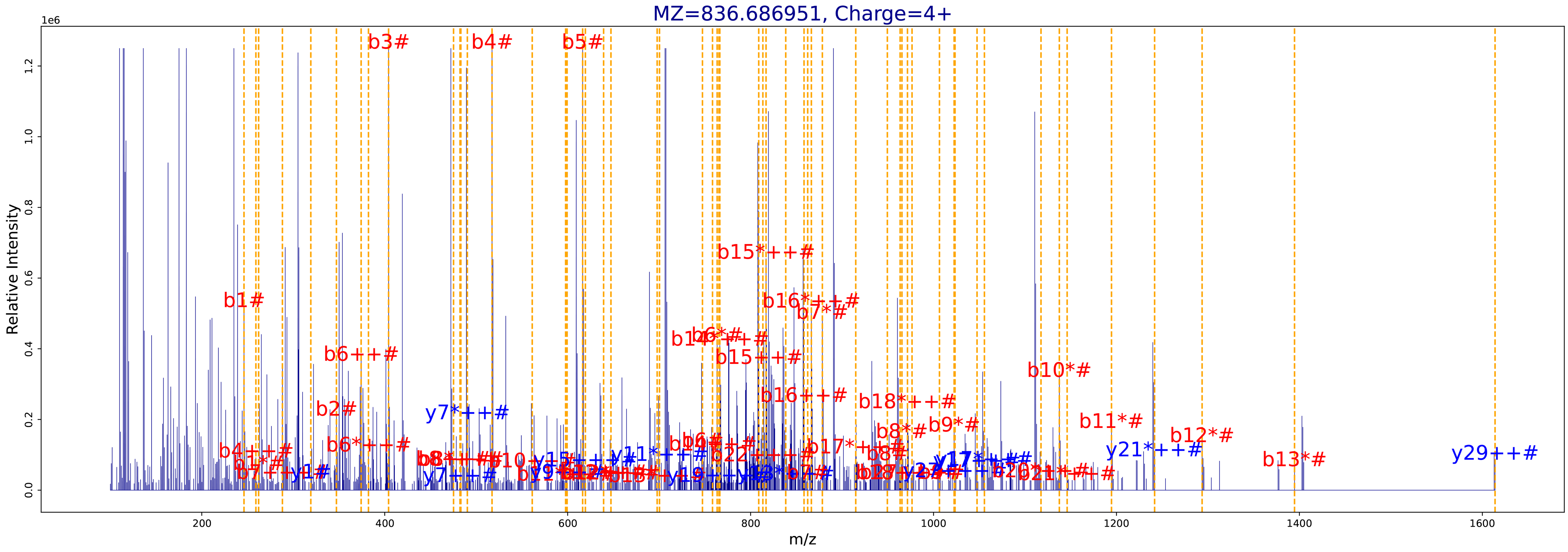

**B**

| | SCAN INFO |
|---|---|
| Raw | JAL_NOa2_iTR_Fr1 |
| Scan | 108616 |
| Charge | 3 |
| RT | 368.4 |
| DeltaM | 15.995138 |
| Label | iTRAQ4plex, Carbamidomethyl, Carbamidomethyl, iTRAQ4plex |
| MH | 3890.926731 |
| E-score | 5.969404 |

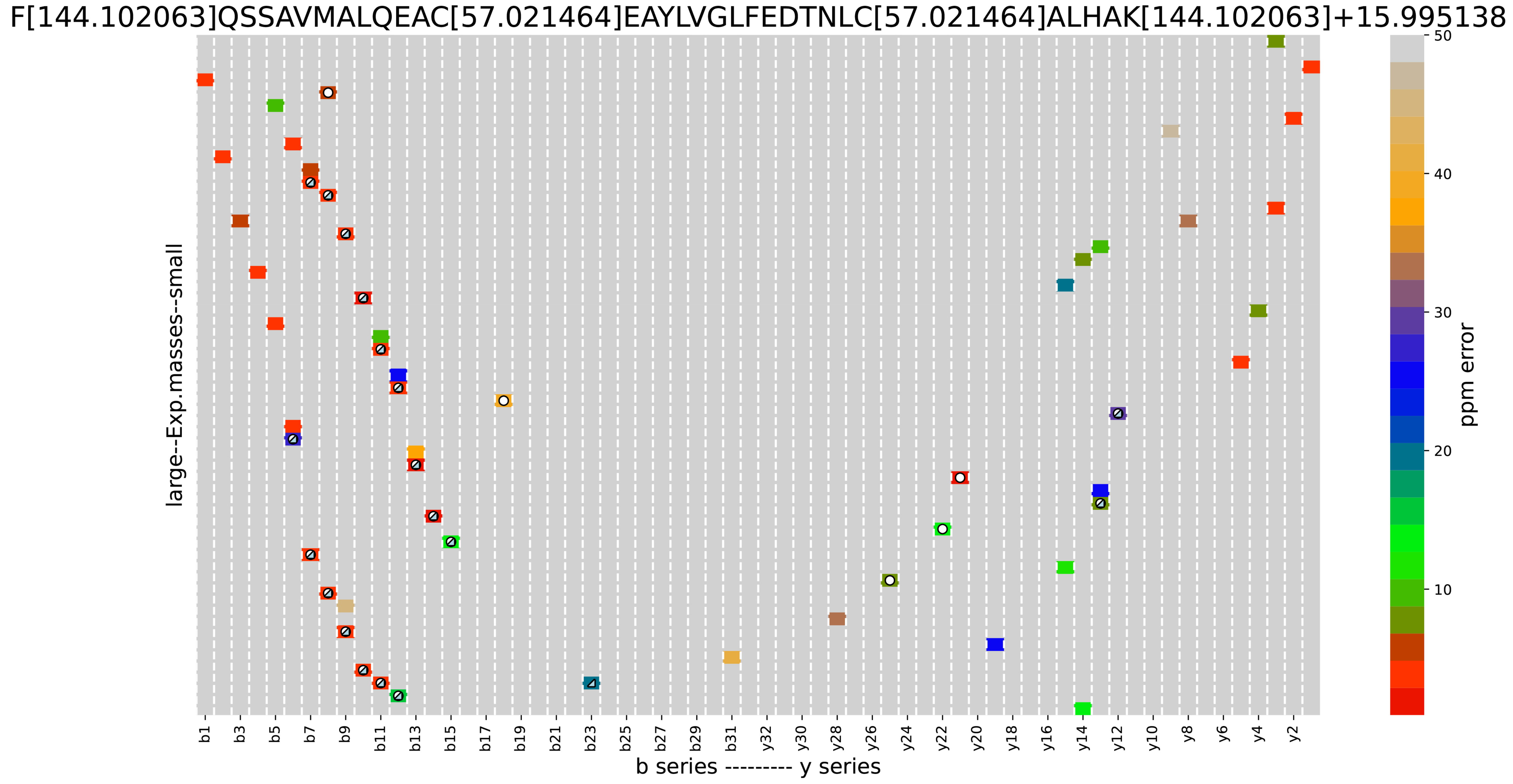


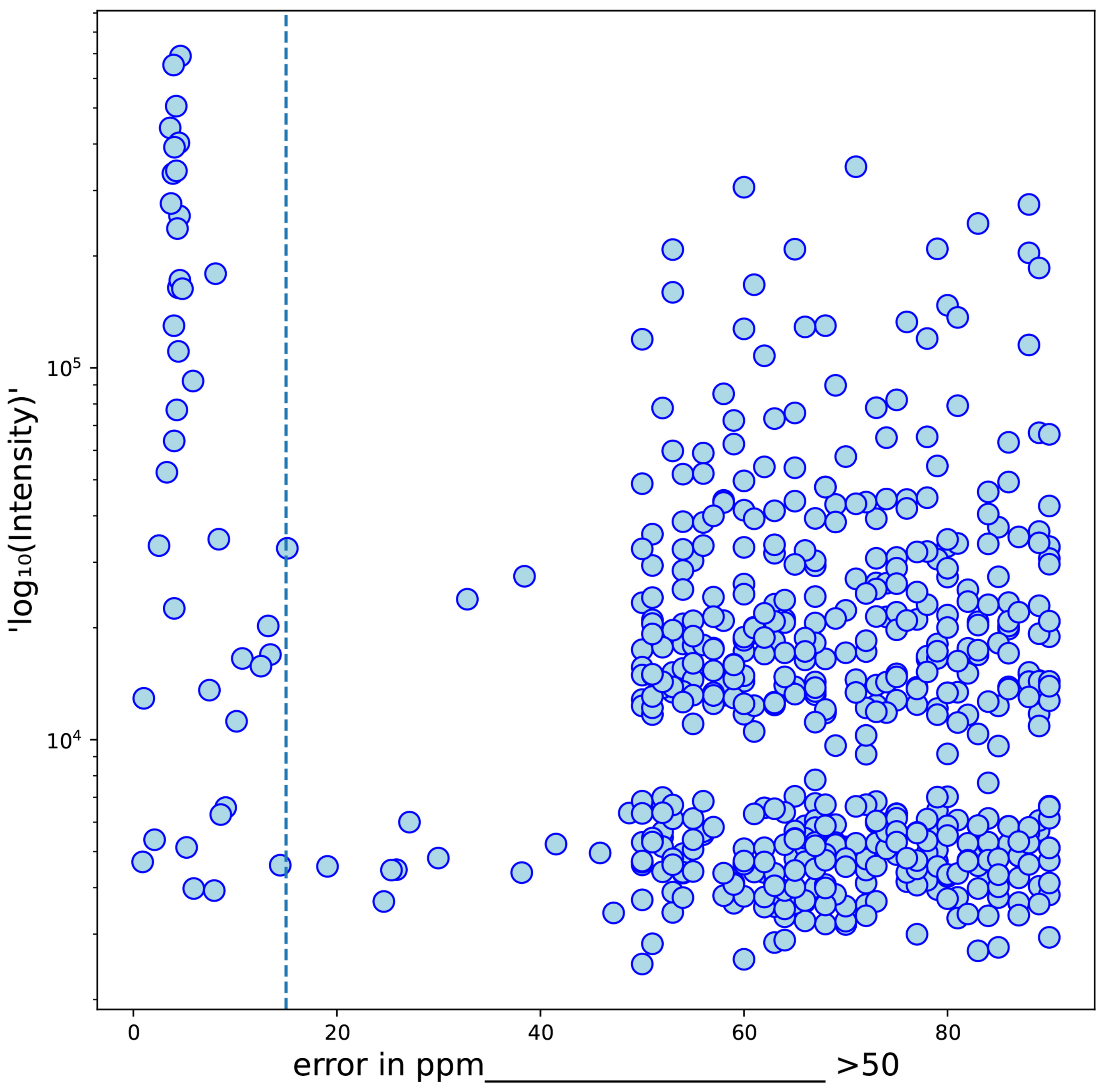


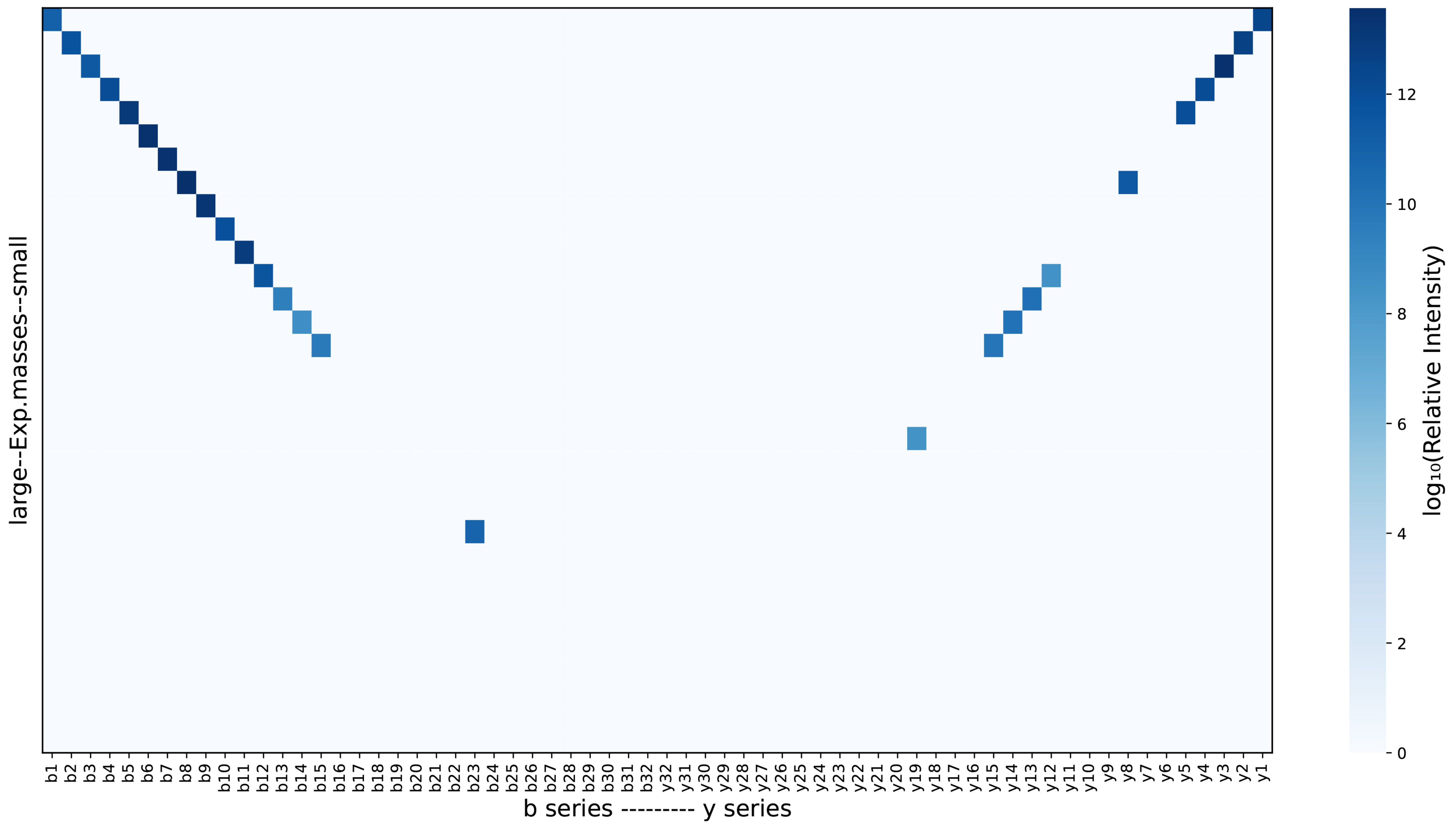


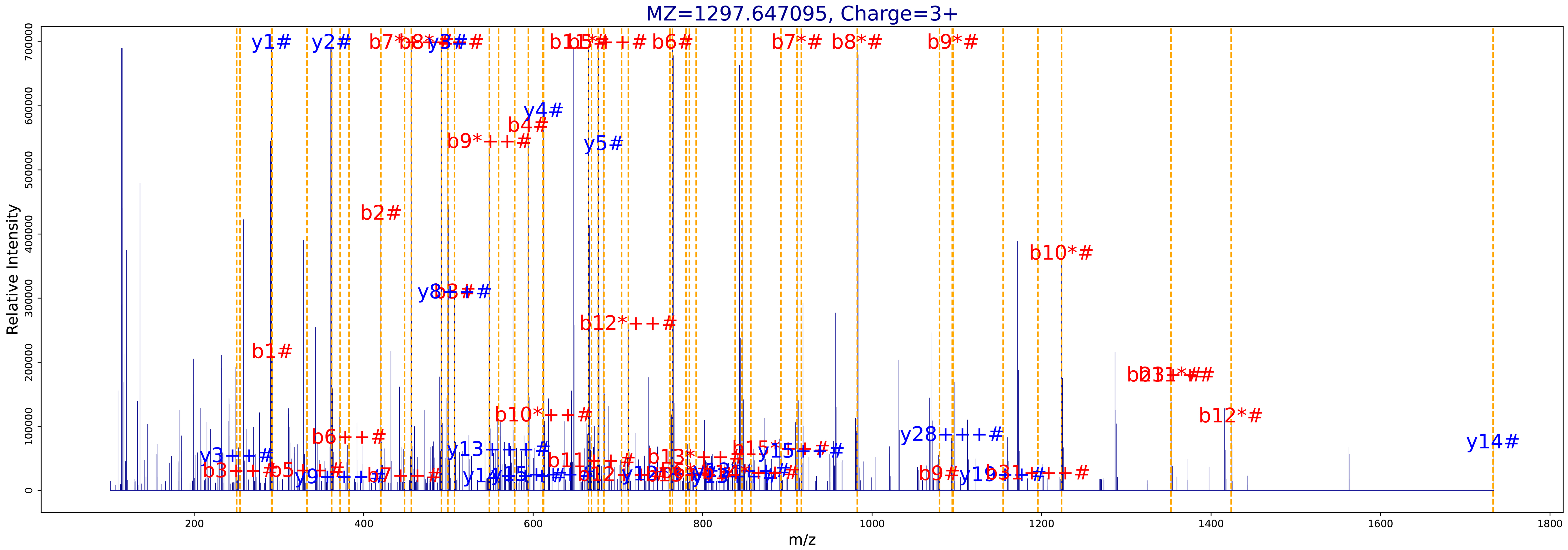

**C**

| | SCAN INFO |
|---|---|
| Raw | JAL_NOa2_iTR_Fr1 |
| Scan | 63376 |
| Charge | 4 |
| RT | 197.86 |
| DeltaM | 15.995138 |
| Label | iTRAQ4plex |
| MH | 3341.72524 |
| E-score | 72.442622 |

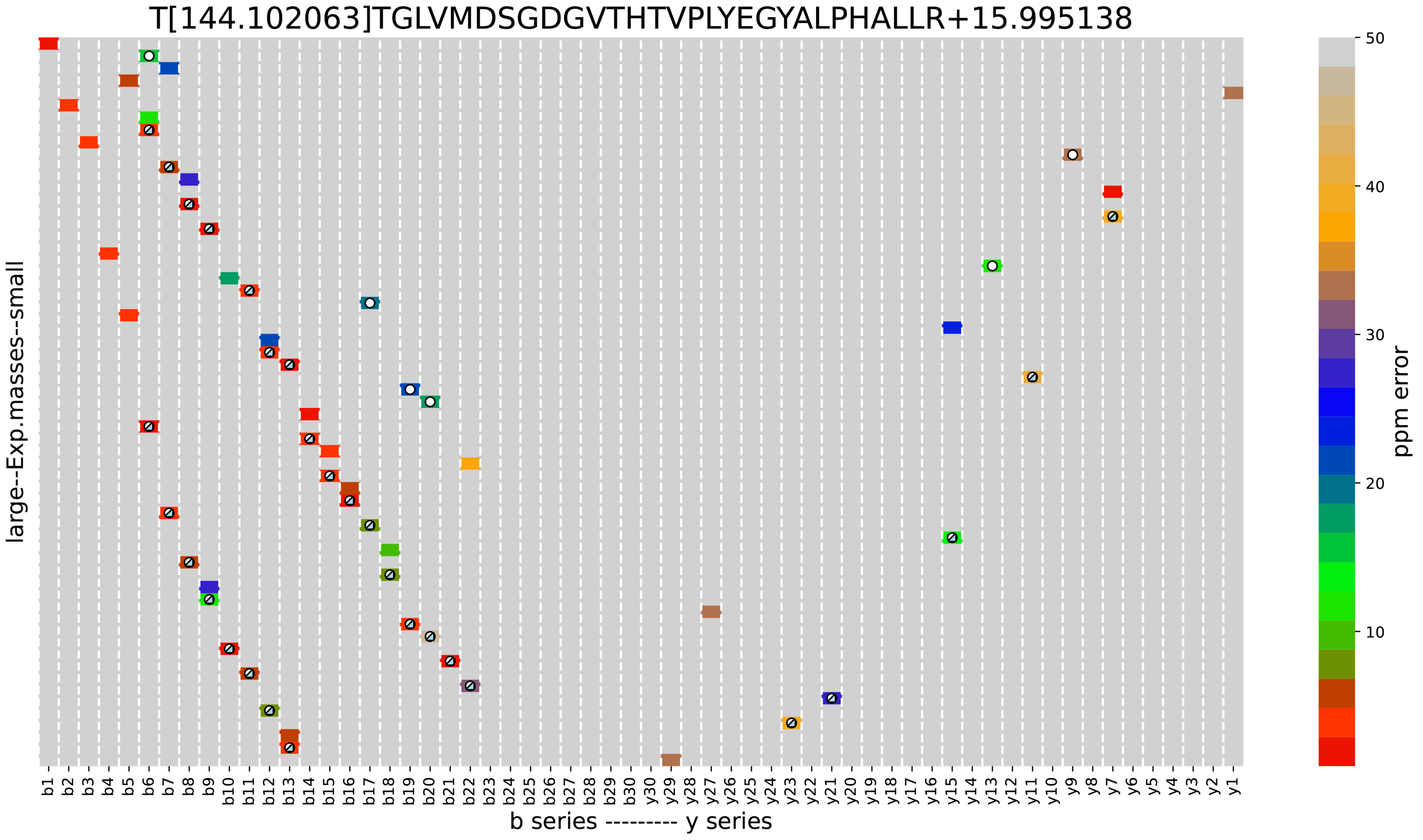


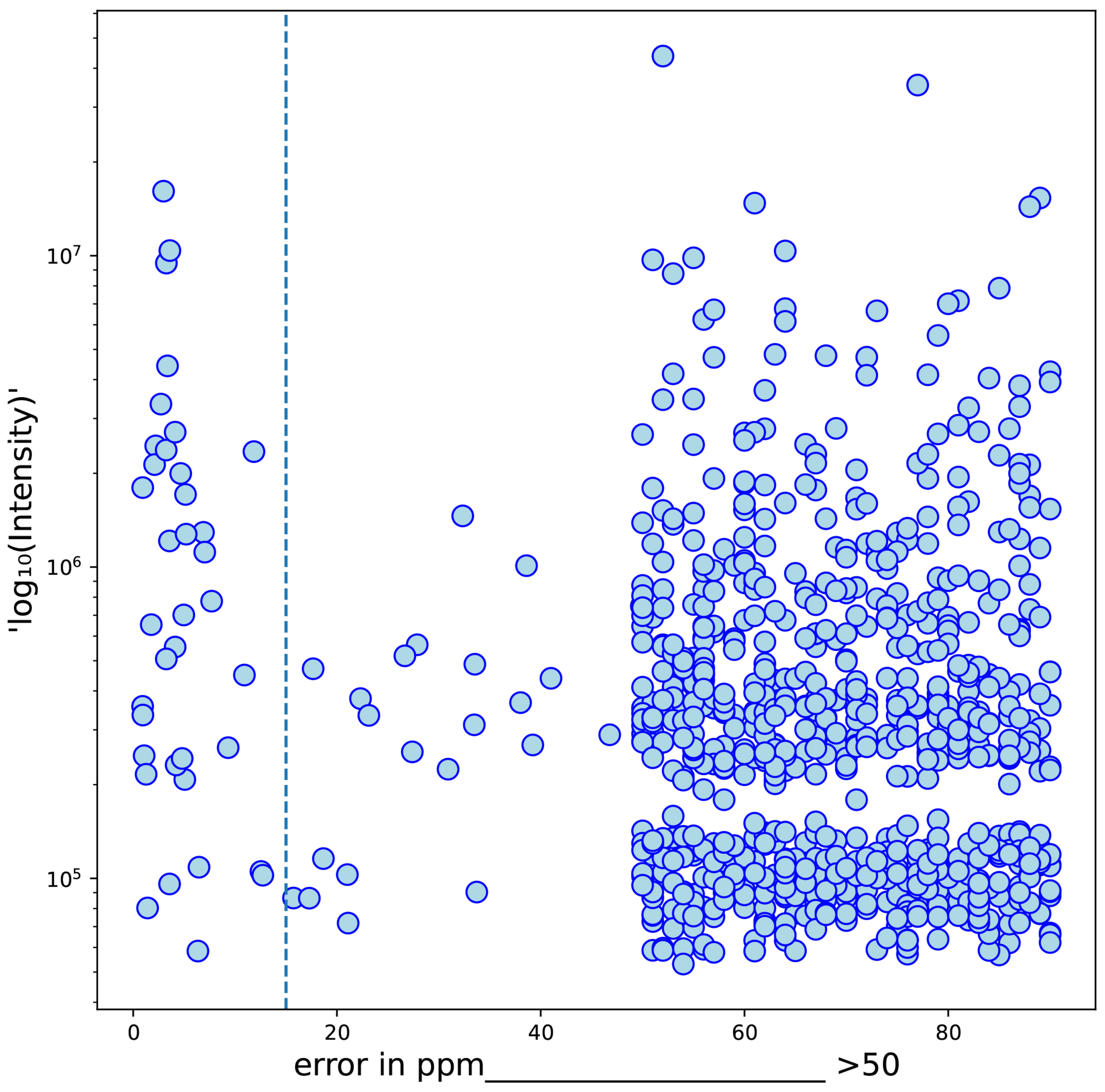


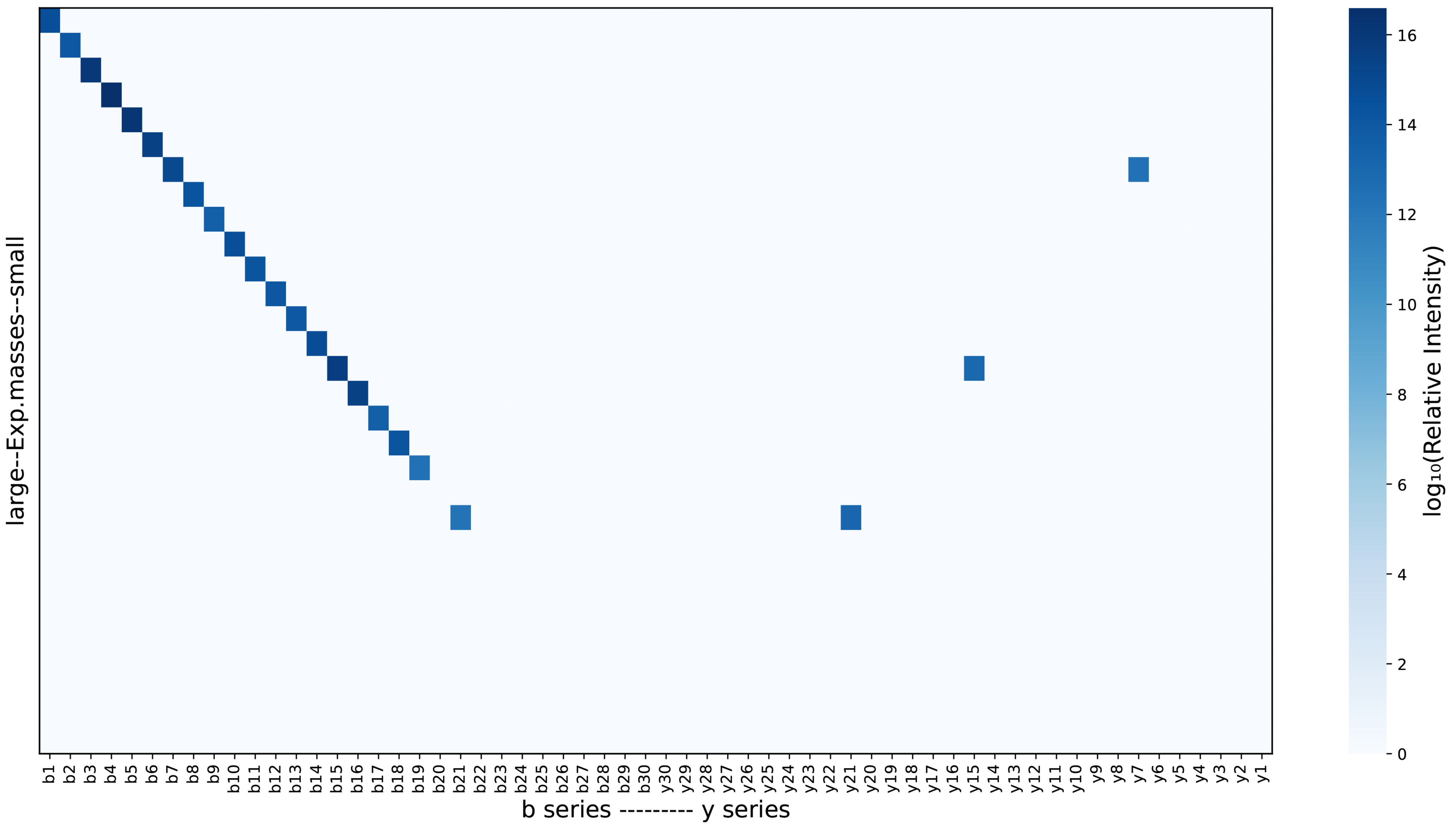


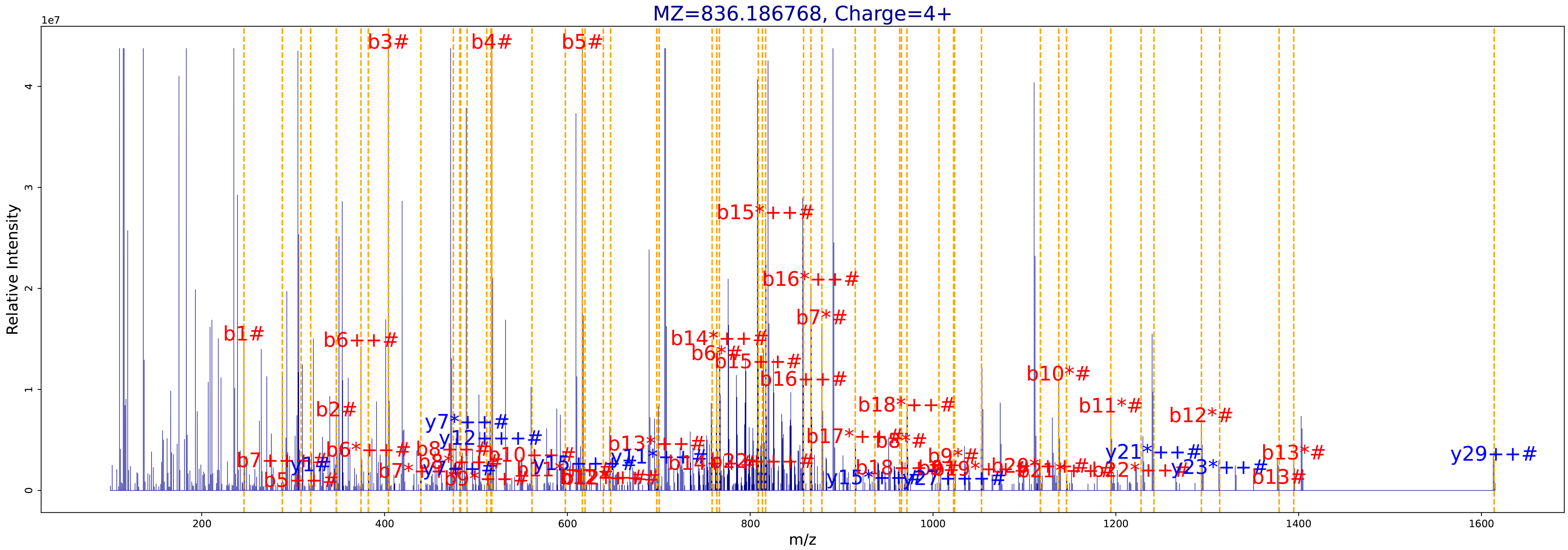

**D**

| SCAN INFO | |
|---|---|
| Raw | JAL_NOa2_iTR_Fr1 |
| Scan | 91769 |
| Charge | 4 |
| RT | 302.54 |
| DeltaM | 31.989925 |
| Label | iTRAQ4plex, iTRAQ4plex, iTRAQ4plex |
| MH | 3481.800436 |
| E-score | 0.628162 |

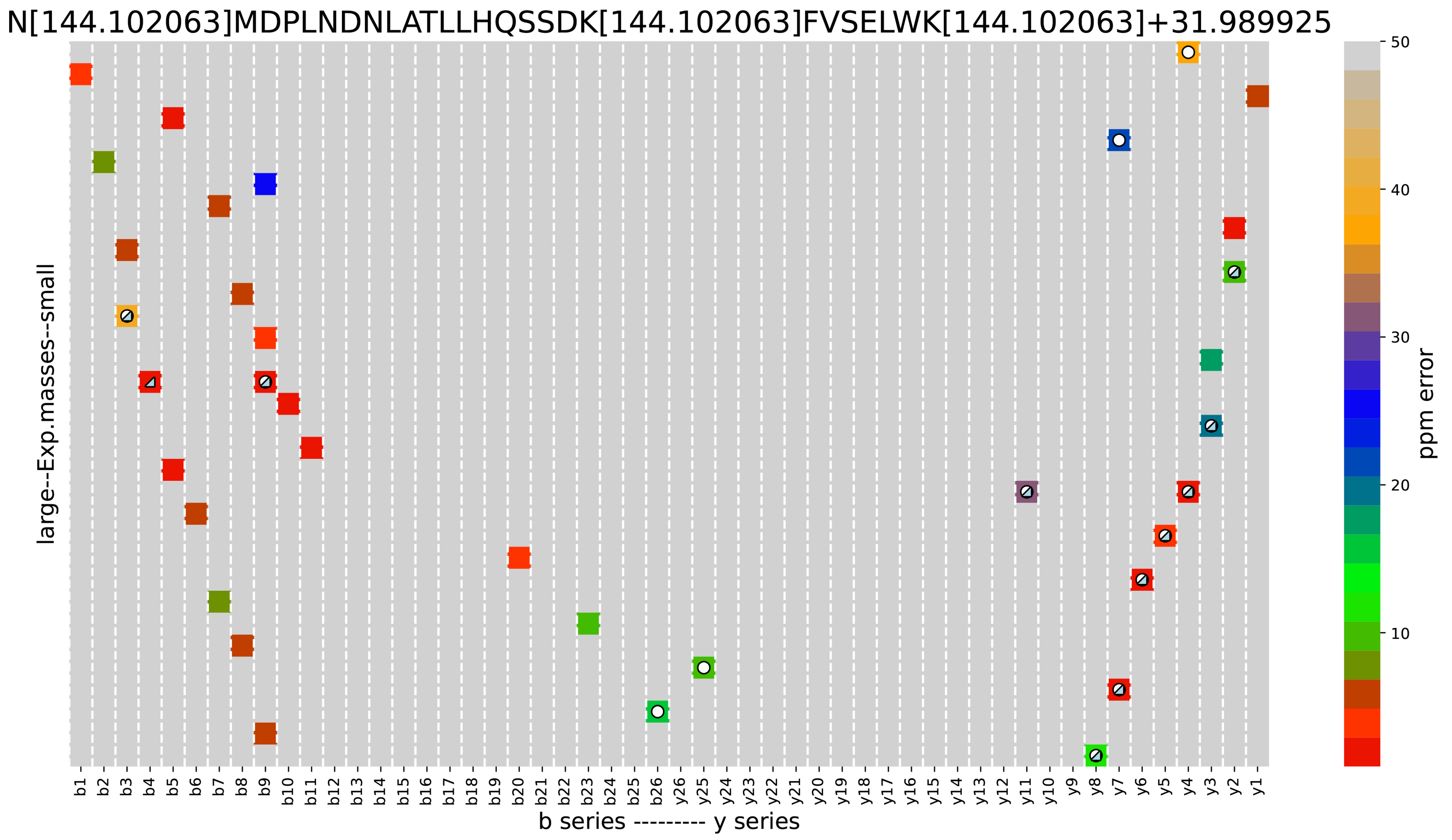


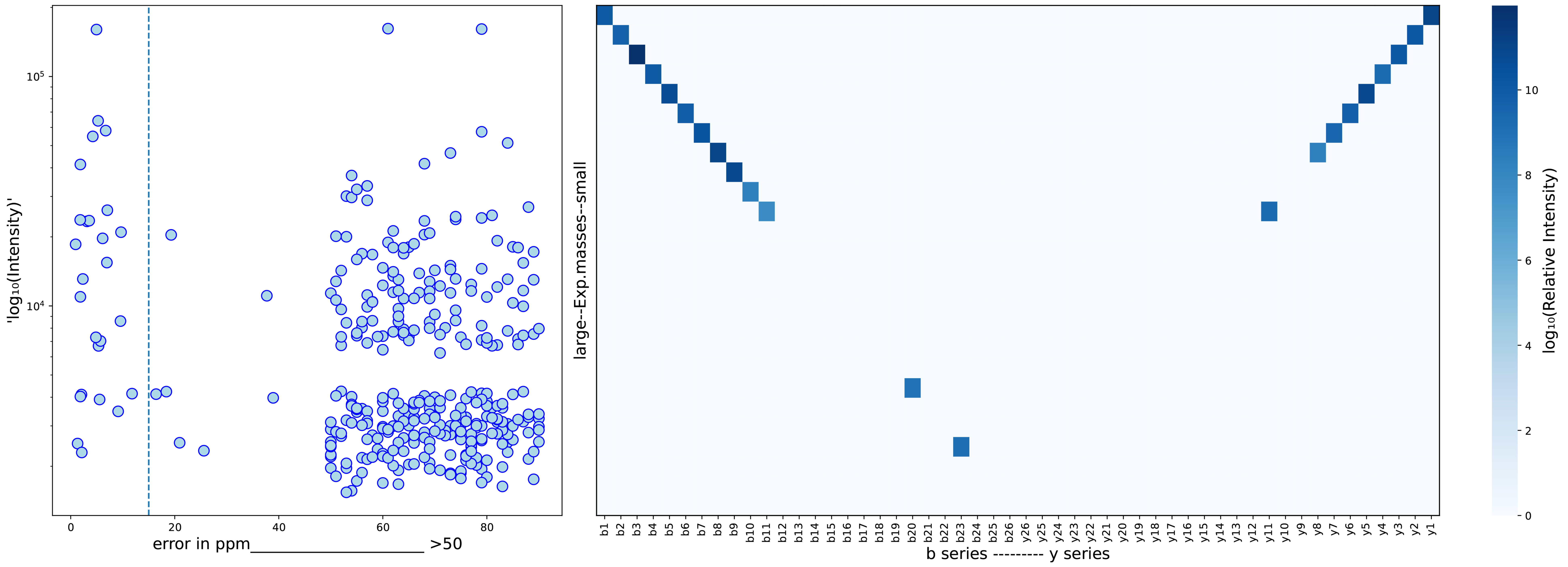


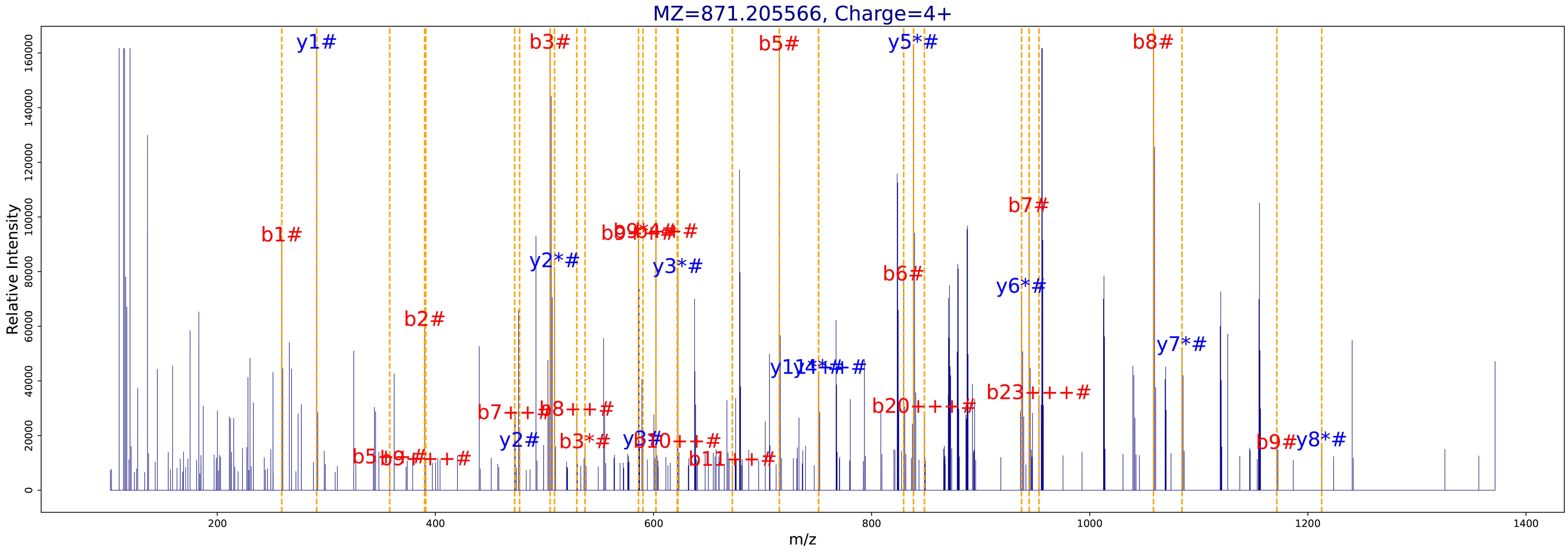

**E**

| SCAN INFO | |
|---|---|
| Raw | JAL_NOa2_iTR_Fr1 |
| Scan | 84003 |
| Charge | 4 |
| RT | 272.94 |
| DeltaM | 15.995138 |
| Label | iTRAQ4plex, iTRAQ4plex, iTRAQ4plex |
| MH | 3596.824362 |
| E-score | 4.131782 |

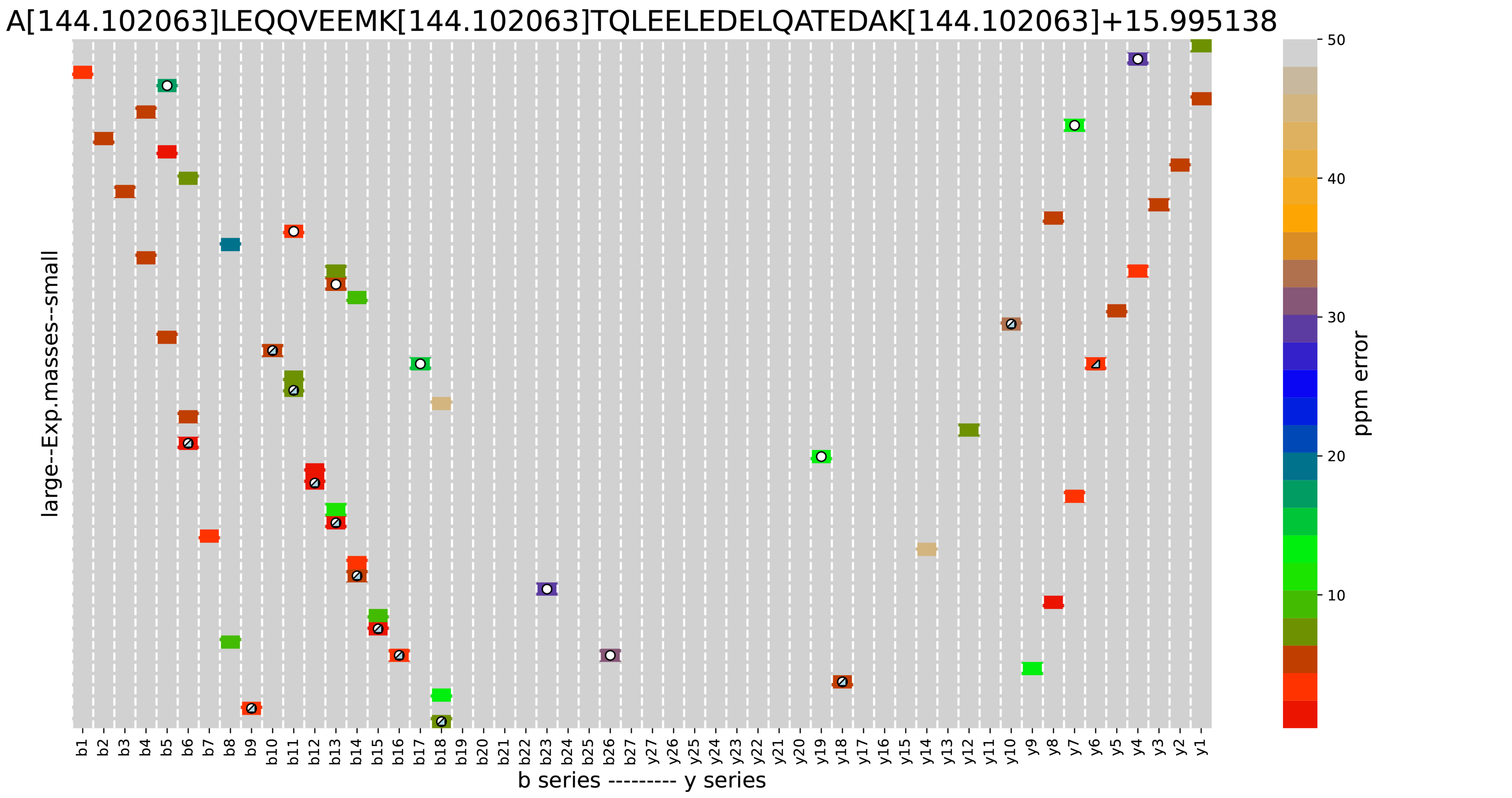


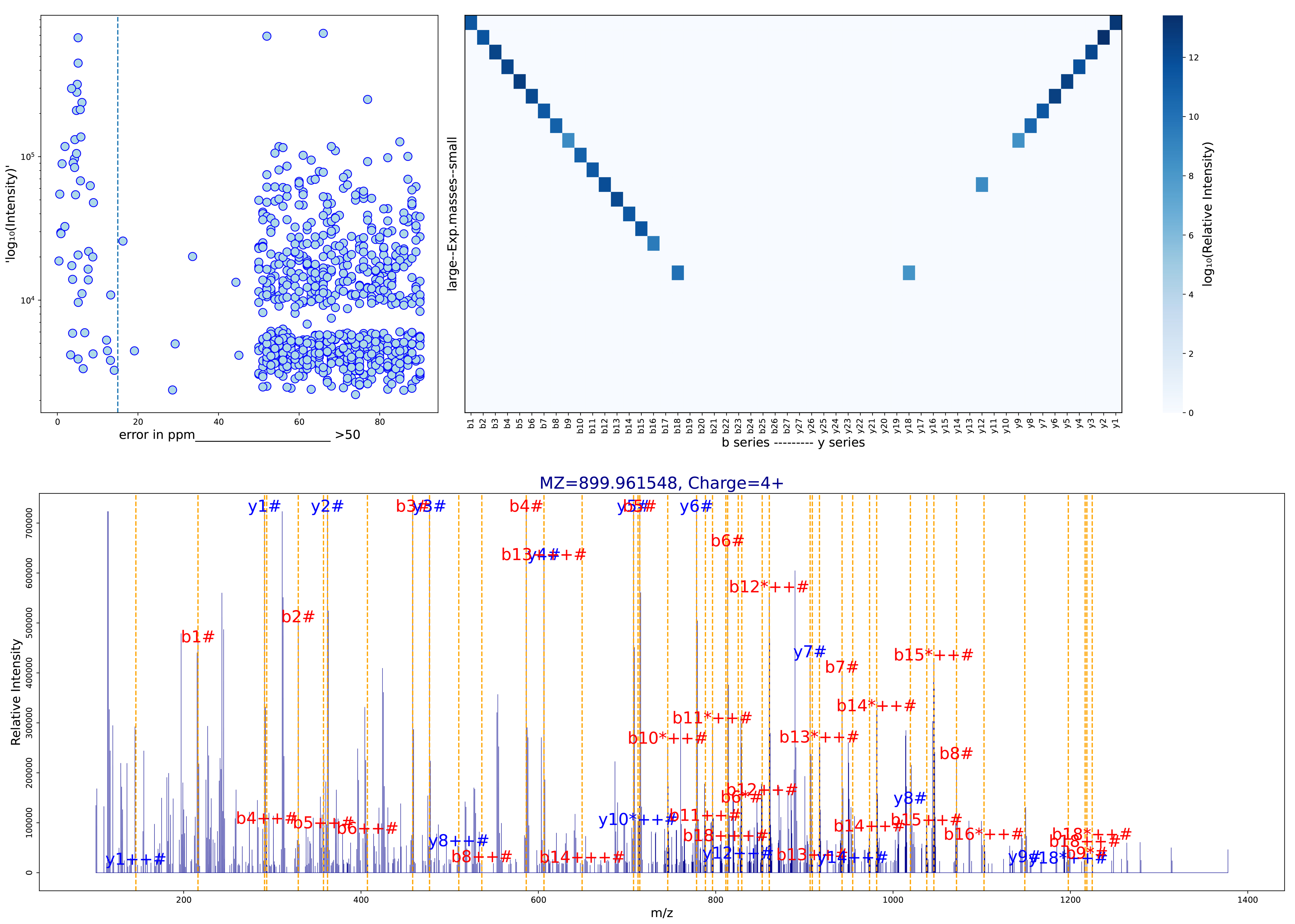

Supplemental Figure 2. Analysis of spectra where only Comet-ReCom made an identification at FDR < 1%, using Vseq. The meaning of each panel is as described in Supp. Fig. 2. These examples correspond to oxidations on M (A, B), on D (C), on K (E) or dioxidation on W (D). All examples show a large number of matched fragments under 15 ppm and high sequence coverage along the b fragment series. Examples B, D and E also show high coverage along the y series.
